# Ultrasound-mediated neuromodulator delivery reveals serotonergic control of context-dependent motivation in primates

**DOI:** 10.64898/2026.05.11.724324

**Authors:** Samuel Winiarski, Mikaella Ty Ngo, William Reith, Marie Habart, Urs Schuffelgen, Mohamed Tachrount, William T. Clarke, William W. Watts, Daniel J. Whitcomb, Trevor Sharp, Eleanor Stride, Matthew FS. Rushworth, Nima Khalighinejad

## Abstract

Causal manipulation of neuromodulatory systems has been constrained by a trade-off between spatial precision and translational relevance: circuit-specific techniques are invasive, whereas systemic pharmacology lacks anatomical specificity. Here, we establish ultrasound-mediated neuromodulator delivery as a non-invasive, spatially precise approach for causal neuropharmacology in awake, behaving primates. By transiently opening the blood–brain barrier, we delivered systemically administered serotonin to the perigenual anterior cingulate cortex (pgACC) during a decision-making task. Combining computational modelling with multimodal imaging, pupillometry and histology, we show that elevating pgACC serotonin altered local neural activity and weakened the influence of reward environment on motivational state while leaving sensitivity to immediate reward opportunities intact. This was accompanied by reduced functional coupling, decreased local glutamatergic metabolite signal and increased serotonergic signal at the target, with repeated delivery well tolerated. Together, these findings establish a translatable platform for focal neuropharmacology and identify pgACC serotonin as a regulator of context-dependent motivation.

## Main Text

Serotonin (5-hydroxytryptamine; 5-HT) fundamentally shapes brain function and behaviour. Clinically, serotonin is the primary pharmacological target for a host of psychiatric disorders, yet its role in cognition remains poorly understood. Some previous findings have demonstrated serotonin’s role in tracking environmental statistics (1–5), whereas others have implicated serotonin in changes of behavioural strategy, such as between impulsivity and persistence or exploration and exploitation (6–11).

To fully disentangle the cognitive function of serotonin, causal methods are required that are both spatially precise and translatable across species, enabling manipulation of defined neural circuits while remaining clinically relevant. Existing approaches operate at disparate spatial scales: systemic pharmacology is clinically relevant but anatomically diffuse, whereas circuit-specific manipulations are typically invasive and difficult to translate to primates, including humans. This trade-off has limited the ability to causally link neuromodulators to defined brain circuits and cognitive functions in species most relevant to human neuropsychiatric disease. Here, we introduce a framework that resolves this trade-off by non-invasively delivering serotonin to a defined brain region, without genetic or surgical manipulation, to reveal its causal influence on neural activity and behaviour.

Low intensity transcranial ultrasound (TUS), when combined with phospholipid-coated microbubbles (MB), can transiently and focally open the blood-brain barrier (BBB) via mechanical forces generated by cavitation at the ultrasound focus (Fig. 1C) (12). Because the BBB restricts molecular transport between blood and brain, this approach allows for spatially targeted delivery of molecules that would otherwise not enter the brain. Ultrasound-mediated BBB opening has been shown to be safe and effective in non-human primates (13–16), raising the possibility that it could serve as a general platform for focal neuropharmacological manipulation during behaviour.

**Figure 1.**
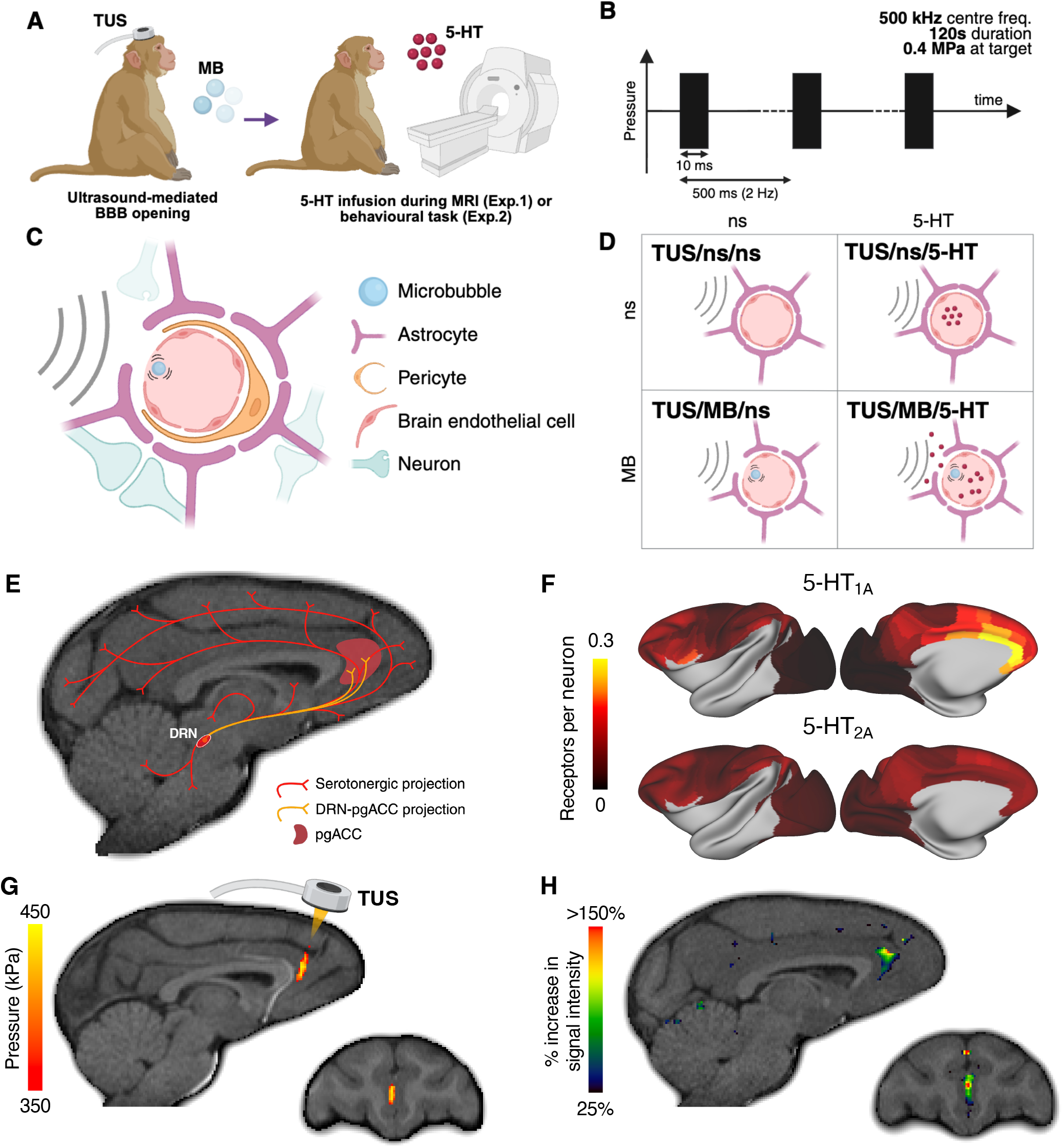
Experiment overview and validation of focal BBB opening at the pgACC. (A) Experimental overview. TUS combined with MB is used to open the BBB at the pgACC. After BBB opening, serotonin is continually infused during MRI scans, including resting-state functional MRI and MRS in Experiment 1, during behavioural testing in Experiment 2, and prior to postmortem histology in Experiment 3. (B) TUS parameters used for BBB opening: 500 kHz centre frequency, 10 ms pulse length, 500 ms period, 2 Hz pulse repetition frequency, 2 % duty cycle, 120 s duration, yielding a peak negative pressure of 0.4 MPa at the target. (C) Schematic showing ultrasound-induced MB cavitation in a cerebral blood vessel, and subsequent transient opening of the BBB. (D) Experimental conditions. TUS/ns/ns: TUS without MBs, followed by normal saline (ns) infusion; TUS/ns/5-HT: TUS without MBs, followed by serotonin (5-HT) infusion; TUS/MB/ns: TUS with MBs followed by ns infusion; TUS/MB/5-HT: TUS with MBs, followed by 5-HT infusion. (E) Schematic illustrating serotonergic projections to cortical and subcortical regions in a macaque brain. The pgACC is highlighted in red, and projections from the DRN to the pgACC are shown in yellow. (F) Cortical distribution of 5-HT_1A_ and 5-HT_2A_ receptors in macaque shown on the cortical surface. Data for receptor densities from (27). pgACC has high density of inhibitory 5-HT1A receptors. The colour bar shows the density of receptors per neuron. (G) Subject-specific ultrasound simulation used to confirm transducer positioning and estimate acoustic pressure at the target. The simulated pressure field is masked to the brain and thresholded at 0.35 MPa. Representative transducer position is shown for clarity. (H) Representative example of voxel-wise signal-intensity ratio maps showing contrast enhancement within a whole-brain grey-matter mask, computed as ratio between a BBB opening session and a corresponding non-BBB opening session acquired within the same experimental round (for data from all sessions see Table S2). Graphics were created with BioRender.

Here, by transiently opening the BBB, we deliver systemically administered serotonin to a defined cortical target and assess the consequences across multiple levels of analysis, including histology, magnetic resonance spectroscopy (MRS), resting-state functional MRI (rs-fMRI), computational behavioural modelling, and pupillometry. In doing so, we establish a translatable framework for causal neuropharmacology in awake, behaving primates and use it to address a longstanding question about serotonergic function.

We chose serotonin as a biological demonstration of the platform because it plays a central role in motivational control. We recently showed that the dorsal raphe nucleus (DRN), the principal serotonergic nucleus of the brain, tracks the average value of the environment and regulates transitions between latent motivational states in macaques (17), reconciling two prevailing views of serotonergic function: tracking reward-related features of an animal’s environment (1–5) and controlling changes in an animal’s behavioural strategy (6–11). However, although predominantly serotonergic, the DRN contains multiple other neuronal populations, including glutamatergic, dopaminergic and GABAergic cells (18,19). Moreover, ascending serotonergic projections from the DRN influence behaviour through diverse cortical targets with distinct 5-HT receptor profiles. Consequently, it remains unresolved how serotonergic signalling downstream of the DRN – particularly within frontal cortical circuits – contributes to motivational state control.

One major cortical target of the DRN is the perigenual anterior cingulate cortex (pgACC), a region rich in inhibitory 5-HT_1A_ receptors (20). Converging evidence from studies using imaging, lesion, and electrophysiological approaches implicates the pgACC in value-based decision-making, integration of reward and contextual information, and in mediating the influence of internal motivational state on behaviour (21–24). We therefore test the hypothesis that serotonergic signalling within the pgACC mediates part of the DRN’s influence on motivational state. Untangling how serotonergic signalling within frontal cortical circuits shapes motivation is critical for understanding serotonin’s role in clinical conditions characterised by disrupted motivation, including depression and anxiety.

Using ultrasound-mediated neuromodulator delivery, we show that focal serotonin delivery to the pgACC attenuates local glutamatergic metabolite signal, suppresses functional coupling between pgACC and distributed frontal networks, and weakens the influence of the reward environment on motivational state while leaving sensitivity to immediate reward opportunities unchanged. We also demonstrate that repeated BBB opening procedures are safe and well-tolerated in non-human primates over a period of months, positioning TUS-mediated neuromodulator delivery as a tool for probing brain circuits at the mesoscale. Together, these findings identify pgACC serotonin as a regulator of context-dependent motivation and establish ultrasound-mediated neuromodulator delivery as a non-invasive, spatially precise and translatable platform for causal neuropharmacology in awake, behaving primates, with direct relevance for understanding and targeting circuit-level dysfunction in disorders of motivation.

### Ultrasound-mediated BBB opening enables focal serotonin delivery to the pgACC

In this study, we use ultrasound-mediated neuromodulator delivery to causally manipulate serotonergic circuitry in rhesus macaques (*Macaca mulatta*) by delivering serotonin selectively to the pgACC (Fig. 1A). Low-intensity TUS was applied at the pgACC (0.4MPa in situ; see Fig. 1B for TUS parameters) following intravenous MB injection, to induce focal, transient BBB opening via cavitation within the cerebral vasculature (Fig. 1C). This technique generates localised mechanical forces at the blood vessel wall, resulting in a reversible increase in BBB permeability (mean size of permeabilised region = 87 mm^3^). Following BBB opening, serotonin was infused systemically via the saphenous vein (600 ng/kg/min; see Fig. S1 for dose determination). Because serotonin does not cross an intact BBB, diffusion into the brain was restricted to the sonicated pgACC, enabling focal serotonin delivery.

To isolate the effects of focal serotonin delivery to the pgACC, we used a 2×2 factorial design implemented across two experiments (Experiments 1&2), independently manipulating MB administration (on/off) and serotonin infusion (on/off) (Fig. 1D & Table S1). TUS was applied in all four conditions. This design dissociated the effects of focal serotonin delivery to pgACC from those of systemic serotonin infusion or BBB opening alone, while controlling for any potential neuromodulatory effects of TUS. In a third experiment (Experiment 3), following repeated BBB opening over several months, we performed histological analyses to both assess tissue integrity and verify the site of serotonin action. The latter used immunostaining for serotonin to demonstrate detectable serotonin delivery to the pgACC.

In Experiment 1 (n=3), we examined the effects of focal serotonin delivery to the pgACC on brain chemistry and functional connectivity using single voxel magnetic resonance spectroscopy (1H-MRS) and resting-state functional magnetic resonance imaging (rs-fMRI). In Experiment 2 (n=2), we investigated the behavioural and autonomic physiological effects of focal serotonin manipulation of the pgACC using a previously described motivated decision-making task and pupil size measurements, respectively (17). In Experiment 3 (n=3), we investigated the effects of repeated BBB opening on brain tissue integrity using haematoxylin and eosin (H&E) and Cresyl violet (CV) staining. Further, we assessed serotonin delivery to the pgACC using immunostaining with an anti-serotonin (5-HT) antibody, comparing focal serotonin delivery (n=2) with BBB opening alone (n=1).

We targeted the pgACC for three reasons. First, it receives a direct serotonergic projection from the DRN, through which serotonin modulates neural activity via multiple receptor subtypes including 5-HT_1A_ and 5-HT_2A_ (25) (Fig. 1E). Second, the pgACC has a particularly high density of inhibitory 5-HT_1A_ receptors (20) (Fig. 1F). The expression gradient of 5-HT_1A_ receptors is steep, with a rostrocaudal expression gradient that peaks at or near the pgACC, making this a strong candidate region in which DRN output might exert major influence. Third, converging evidence implicates the pgACC in value-based decision-making and context integration, supporting a role in integrating contextual information from the environment with internal motivational state to influence adaptive behaviour (21–23). Together, these led us to hypothesize that the DRN exerts its control over transitions in motivational state through its serotonergic projections to the pgACC.

We first demonstrated that BBB opening in awake macaques could be achieved focally at the pgACC. We began by performing subject-specific acoustic simulations to determine the TUS stimulation parameters and transducer positioning required to achieve effective targeting of the pgACC. Using k-wave simulations based on skull models derived from Black Bone MRI (26), we identified parameters that yielded a peak negative pressure of 0.4 MPa (corresponding to a spatial peak pulse-average intensity [I_SPPA_] of ~5.3 W/cm^2^) at the pgACC (Fig. 1G), previously shown to induce safe and transient BBB opening in macaques (13). These simulations indicated that achieving this pressure required a free-field I_SPPA_ of 25 W/cm^2^ for monkey M1, 30 W/cm^2^ for monkey M2, and 25 W/cm^2^ for monkey M3.

We then confirmed BBB opening in vivo by quantifying focal extravasation of a gadolinium-based MR contrast agent into the pgACC. Contrast-enhanced T1-weighted MP2RAGE images acquired 10 minutes after gadobutrol injection were processed to generate voxel-wise signal intensity ratio maps comparing BBB-opening sessions with matched non-opening sessions acquired within the same experimental round (TUS/MB/ns relative to TUS/ns/ns; TUS/MB/5-HT relative to TUS/ns/5-HT), within a subject-specific pgACC mask. BBB opening was evidenced by a local increase in contrast-related signal confined to the pgACC (Fig. 1H & Table S2; see methods).

Control experiments using rodent hippocampal slice electrophysiology demonstrated that the TUS parameters used here did not produce measurable neuromodulatory effects in the absence of microbubbles (Fig. S2).

### Focal serotonin delivery to the pgACC alters functional connectivity

Having demonstrated focal BBB opening at the pgACC (Fig. 1H & Table S2), we next investigated whether localised serotonin delivery to the pgACC would induce specific changes in patterns of brain activity. To do this, we acquired rs-fMRI across all experimental conditions (TUS/ns/ns, TUS/ns/5-HT, TUS/MB/ns and TUS/MB/5-HT) and examined the correlation in neural activity between the pgACC (our seed region) and other brain regions (28).

Whole-brain connectivity maps were generated by correlating BOLD activity in the pgACC with activity across the cortex for each experimental condition (Fig. 2A–D). In addition, regions-of-interest (ROIs) corresponding to brain regions with known anatomical connections with the pgACC were defined *a priori* (Table S3; see methods for details) (29,30). These ROIs were used to create connectivity fingerprints in which the radial distance indicates strength of functional coupling to the pgACC, and each angular step demarcates individual target regions (Fig. 2E).

**Figure 2.**
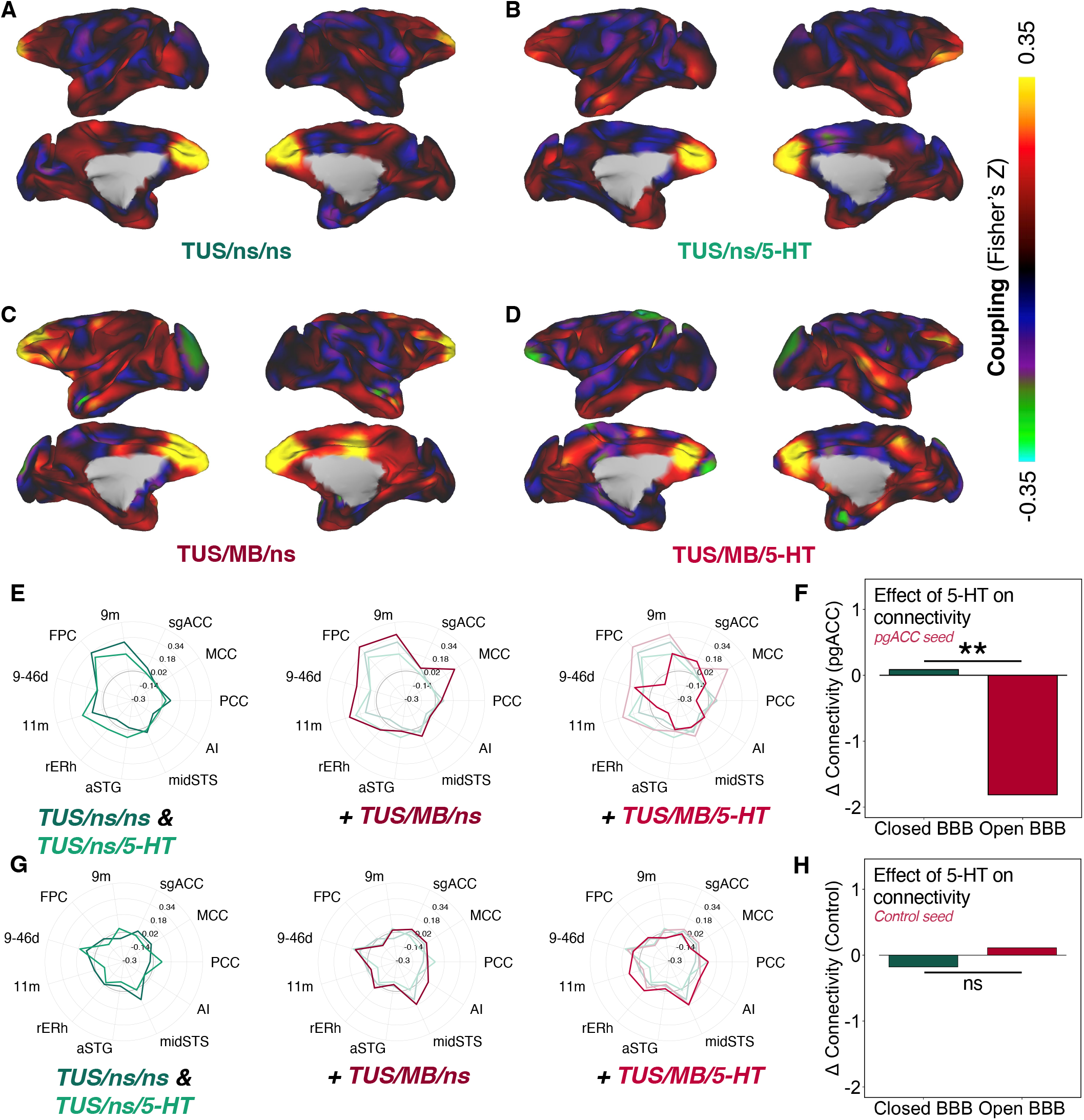
Delivery of 5-HT modulates functional connectivity at the pgACC (n=2). (A-D) Whole brain connectivity maps showing resting-state BOLD correlations between the pgACC seed and all cortical voxels, for each experimental condition. Data were combined across animals within each condition using group-PCA to estimate connectivity patterns at the network level and to capture variance shared across subjects. (E) Connectivity fingerprint plots showing coupling between the pgACC and a priori defined anatomically connected ROIs. Fingerprints are laid out from left to right as TUS/ns/ns and TUS/ns/5-HT, TUS/MB/ns, and TUS/MB/5-HT. The condition of interest is overlaid in brighter colour for comparison. Focal serotonin delivery (TUS/MB/5-HT) suppresses the connectivity pattern compared to BBB opening alone (TUS/MB/ns). (F) Quantification of overall fingerprint connectivity. The net connectivity difference between TUS/MB/5-HT and TUS/MB/ns was found to be significantly larger than that between TUS/ns/5-HT and TUS/ns/ns (Δ_connectivity_ = −1.898, p = 0.00977; non-parametric permutation test). (G) Connectivity fingerprint plots showing coupling between V4d (control region) and ROIs. Fingerprints are laid out from left to right as TUS/ns/ns and TUS/ns/5-HT, TUS/MB/ns, and TUS/MB/5-HT. The condition of interest is overlaid in brighter colour for comparison. Focal serotonin delivery (TUS/MB/5-HT) has no effect on the connectivity pattern compared to BBB opening alone (TUS/MB/ns) at this control seed. (H) Quantification of overall fingerprint connectivity for V4d (control). The net connectivity difference between TUS/MB/5-HT and TUS/MB/ns was not greater than between TUS/ns/5-HT and TUS/ns/ns (Δ_connectivity_ = 0.288, p = 0.514; non-parametric permutation test).

Across conditions, strong functional coupling was observed between the pgACC and adjacent anatomical regions, including the medial prefrontal cortex (mPFC) and orbitofrontal cortex (OFC), as evident in the whole-brain connectivity maps (Fig. 2A–D). Fingerprint analyses further revealed strong coupling between the pgACC and known projection targets, including the frontopolar cortex (FPC), areas 9m, 11m and 9-46d (Fig. 2E). These connectivity profiles were similar in TUS/ns/ns and TUS/ns/5-HT conditions, consistent with the expectation that systemically administered serotonin does not cross an unperturbed BBB.

BBB opening alone (TUS/MB/ns) increased functional connectivity between the pgACC and several anatomically connected regions including the middle cingulate cortex (MCC), FPC, rostral entorhinal cortex (rERh) and areas 9m, 9-46d and 11m (Fig. 2E). Enhancement in functional coupling induced by BBB opening has been previously reported and may arise from transient changes in blood perfusion, oxygenation, or vascular dynamics following BBB permeability alterations (31).

Importantly, focal serotonin delivery to the pgACC (TUS/MB/5-HT) caused marked changes in connectivity patterns relative to BBB opening alone. Specifically, coupling between the pgACC and FPC, MCC, rERh and area 11m was strongly reduced, a pattern opposite to that seen with BBB opening alone. This pattern was evident in both the whole-brain connectivity maps and connectivity fingerprints (Fig. 2D,E).

Quantification of overall fingerprint connectivity revealed a significant difference between TUS/MB/ns and TUS/MB/5-HT conditions (Δ_connectivity_openBBB_ = −1.813, p = 0.00488; see methods). In contrast, no significant difference was observed between TUS/ns/ns and TUS/ns/5-HT conditions (Δ_connectivity_closedBBB_ = 0.085, p = 0.626). Moreover, the net connectivity difference between TUS/MB/5-HT and TUS/MB/ns was found to be significantly larger than between TUS/ns/5-HT and TUS/ns/ns (Δ_connectivity_ = −1.898, p = 0.00977, Fig. 2F). These results suggest that focal serotonin delivery to the pgACC attenuates functional coupling with anatomically connected regions relative to BBB opening alone. These effects were specific to connectivity of pgACC, as no comparable connectivity pattern changes were observed when the seed was placed at a control region (area V4d; Δ_connectivity_openBBB_ = 0.111, p = 0.661; Δ_connectivity_closedBBB_ = −0.178, p = 0.256; Fig. 2G,H). Further, the net connectivity difference for the control seed between TUS/MB/5-HT and TUS/MB/ns was not found to be larger than between TUS/ns/5-HT and TUS/ns/ns (Δ_connectivity_ = 0.288, p = 0.514).

### Focal serotonin delivery to the pgACC reduces local glutamatergic metabolite signal

We then examined whether focal delivery of serotonin to the pgACC produced local metabolic changes in addition to the functional connectivity changes detailed above. We indirectly probed serotonin’s effect by measuring the concentration of metabolites at the pgACC using single-voxel proton magnetic resonance spectroscopy (1H-MRS). This approach enables non-invasive quantification of metabolite concentrations within a defined cortical region in awake macaques.

1H-MRS measures the MR signal arising from multiple metabolites within a single voxel. Each metabolite contributes a characteristic spectral signature and thus allows the observed spectrum to be modelled as a linear combination of the individual metabolite basis functions (Fig. 3A,B). Concentrations were estimated by fitting individual metabolite basis functions to the spectra for each session. In particular, the effect of focal serotonin delivery on glutamate metabolism was of interest, given the role of glutamate as the primary excitatory neurotransmitter in the brain. We first compared glutamate signal to noise ratio (SNR) across conditions to ensure that any observed differences could not be attributed to systemic variation in spectral quality. Glutamate SNR did not vary across conditions (Fig. 3C).

**Figure 3.**
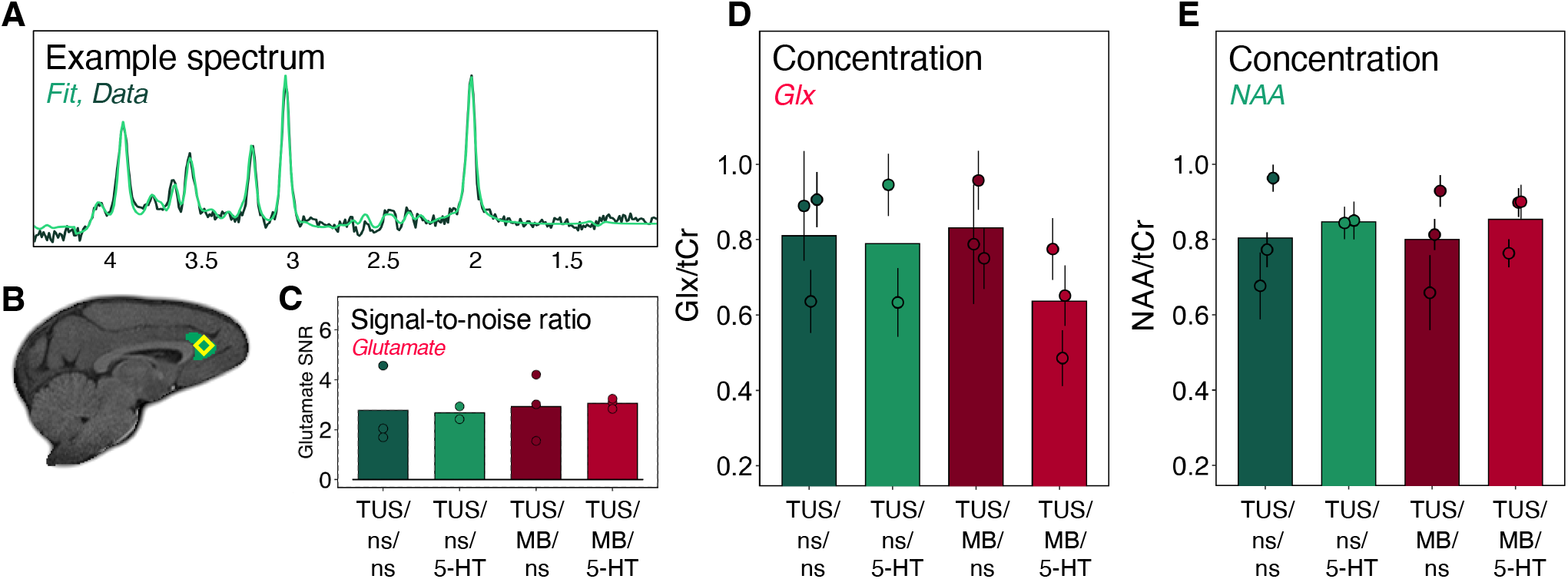
Focal serotonin delivery reduces local glutamatergic metabolite signal at the pgACC (n=3) (A) Example 1H-MRS spectrum showing the raw data (dark) and the fitted model spectrum (light). (B) Characteristic placement of MRS voxel (yellow), shown overlaid on pgACC mask (green). (C) Glutamate signal to noise ratio across conditions, demonstrating comparable spectral quality. Each point represents an individual session. (D) Combined glutamate and glutamine (Glx) concentration estimates relative to total creatine. Each point represents an individual session. Error bars represent Cramér-Rao lower bounds, reflecting uncertainty in metabolite quantification. (E) N-acetylaspartate concentration estimates relative to total creatine signal, also shown with Cramér-Rao lower bounds. One data point is missing from the TUS/ns/5-HT condition due to a technical acquisition error.

Because the glutamate and glutamine signals partially overlap, resulting in correlated concentration estimates, they are included together as a sum in this analysis, denoted Glx. Metabolite concentrations were reported relative to the total creatine (tCr; Cr+PCr) to control for inter-session variability. Error bars indicate Cramér-Rao lower bounds, which reflect the model-estimated uncertainty of metabolite quantification for each individual spectrum.

Relative to BBB opening alone, focal serotonin delivery (TUS/MB/5-HT) was associated with a reduction in Glx levels within the pgACC. In contrast, no change was detected between TUS/ns/ns and TUS/ns/5-HT conditions, consistent with the expectation that systemic serotonin does not cross an intact BBB (Fig. 3D). No comparable changes were observed in the levels of a control metabolite, N-acetylaspartate (NAA), suggesting that the effect was not due to nonspecific tissue alterations (Fig. 3E).

These findings suggest that focal serotonin delivery to pgACC is associated with a reduction in local glutamatergic metabolite signal. Given the high density of inhibitory 5-HT_1A_ receptors in pgACC, this reduction may reflect 5-HT_1A_ mediated suppression of local excitatory circuits.

### Focal serotonin delivery to the pgACC reduces the influence of environment value on reward pursuit

In Experiment 1, we demonstrated that focal serotonin delivery to the pgACC altered functional connectivity and local glutamate metabolism. We next investigated the effects of focal serotonin delivery to the pgACC on behaviour (Experiment 2, n=2). Monkeys performed an established decision-making task in which they encountered sequential reward opportunities within varying reward environments (17).

The behavioural task was conducted inside the MRI scanner following BBB opening. After positioning the animals in the scanner and prior to task onset, contrast-enhanced structural images (MP2RAGE) were acquired following gadobutrol administration to confirm BBB opening (Fig. 1H and Table S2). The task was subsequently performed with continuous serotonin or saline infusion (depending on condition), which began a few minutes before task onset and continued until shortly after task completion. Two animals completed all four conditions in a within-subject design (Fig. 1D), with seven repetitions per condition (total sample per condition: 2 animals x 7 repetitions = 14).

On each trial, animals encountered a reward opportunity defined by magnitude (signalled by offer colour) and reward probability (signalled by number of dots) (Fig. 4A). Animals could choose to pursue a given opportunity, incurring a short time delay before proceeding to the next offer, or let the offer pass and proceed to the next trial without delay. Reward opportunities were encountered within four distinct block types that varied in overall reward environment: Rich-predictable, Rich-stochastic, Poor-predictable, and Poor-stochastic (Fig. 4B,C). Rich blocks had a higher mean reward probability (0.80) than Poor blocks (0.55). Stochasticity was defined by the width and shape of the probability distribution: stochastic blocks followed a broad uniform distribution (SD = 0.13), while predictable blocks followed a narrow normal distribution (SD = 0.05). Each block consisted of 40-50 trials (see Methods for details).

**Figure 4.**
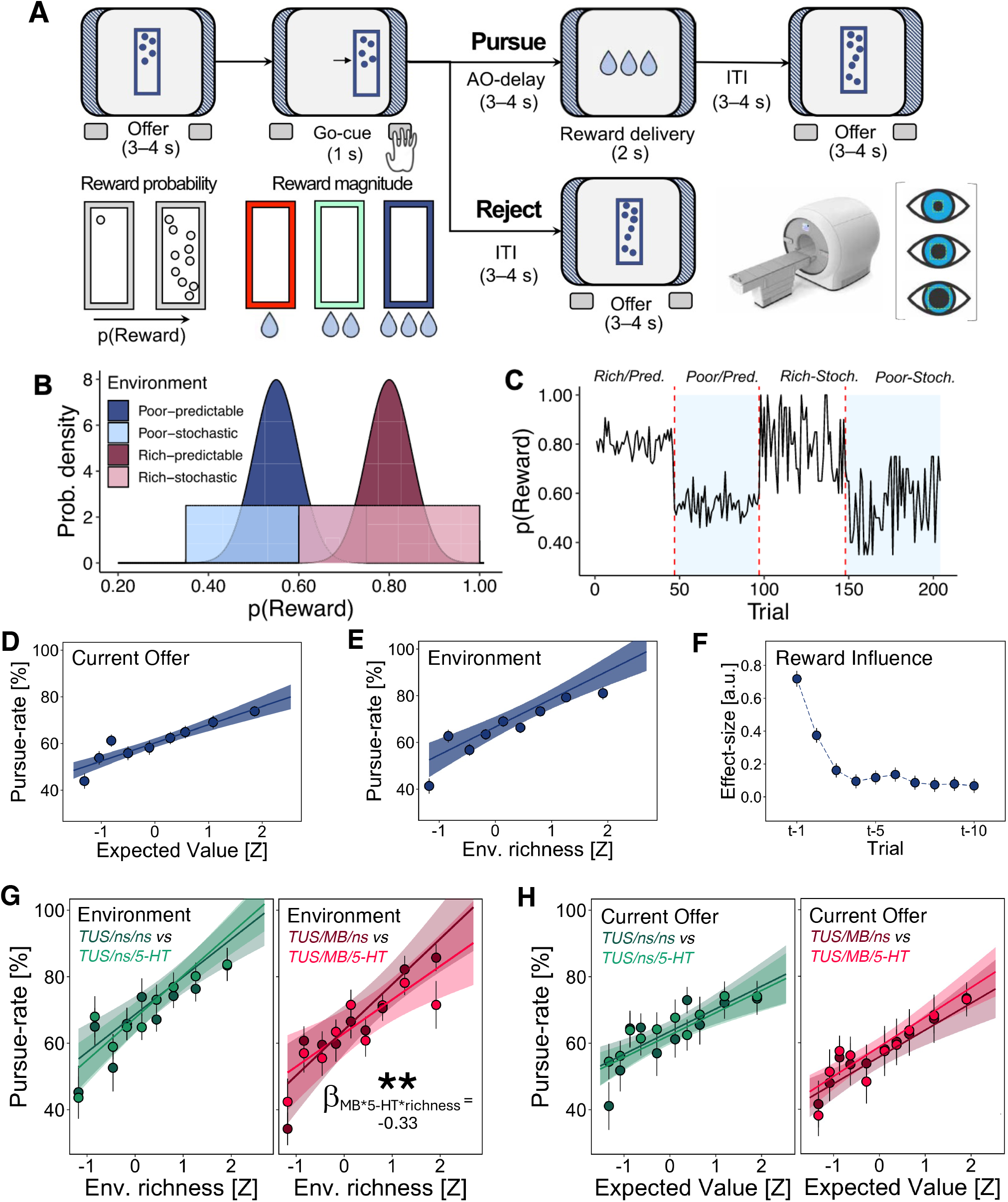
Focal serotonin delivery reduces the effect of environment value on decisions in a pursue-reject task. (A) The behavioural task involved sequential encounters with reward opportunities that varied in reward magnitude (stimulus colour) and reward probability (dots per stimulus). Animals (n=2) performed the task during functional MRI and pupillometry. (B) Probability density of reward offers, across the four block types. Predictable blocks followed a normal distribution (SD = 0.05), while stochastic blocks followed a uniform distribution (SD = 0.13). Rich blocks had a mean p(reward) of 0.8; poor blocks, 0.55. (C) A characteristic depiction of changes in reward probability as a function of reward environment in a single session. Each session comprised four environments covering each permutation of richness and stochasticity. (D) Pursue rate as a function of expected value of the current, specific reward opportunity (reward size x reward probability). Dots and whiskers indicate mean & SEM of pursue rate in deciles of expected value for each animal and session. (E) Pursue rate as a function of environment richness (mean rewards received in the past 5 trials). Dots and whiskers indicate mean & SEM of pursue rate in deciles of environment richness for each animal & session. (F) Effect of reward history window on behaviour. Regression coefficients for environment richness computed over varying window lengths (3–10 trials). Consistent with previous work, a five-trial window was used for analyses; results were unchanged across alternative window lengths. (G) Focal serotonin delivery (TUS/MB/5-HT) significantly attenuated the effect of environment richness on pursue-rate (GLM2.2; β_MB*5-HT*Env.richness_ = −0.33, p = 0.00116). Dots and whiskers indicate mean pursue-rate ± SEM within deciles of environment richness (H) Focal serotonin delivery (TUS/MB/5-HT) did not change the effect of expected value of the current offer on pursue-rate (GLM2.1; β_MB*5-HT*expected-value_ = 0.053, SE = 0.09, p = 0.550). Dots and whiskers indicate mean pursue-rate ± SEM within deciles of expected value.

Mixed effect binomial GLMs were used to quantify the effect of various predictors on pursue-reject decisions (see Methods). First, we looked at the impact of each individual reward opportunity (the current offer) on behaviour. Animals were more likely to pursue reward opportunities with higher reward magnitude and probability (GLM1.1; β_reward-magnitude_ = 0.23, SE = 0.03, p = 1.67e-14; β_reward-probability_ = 0.27, SE = 0.12, p = 0.0317), and with higher expected value (reward probability x reward magnitude) (GLM1.2; β_expected-value_ = 0.33, SE = 0.03, p < 2e-16; Fig. 4D). Effects remained significant when models were fit separately for each animal (Table S4).

Next, we examined the influence of the reward environment, within which a given opportunity arose, on behaviour. Consistent with previous work (17), recent reward history influenced behaviour. We operationalised environmental value (“environment richness”) as the mean reward obtained over the preceding five trials (Fig. 4F). Animals were more likely to pursue opportunities when recent reward history was high, i.e. when the recent environment was rich (GLM1.3; β_Env.richness_ = 0.75, SE = 0.13, p = 2.18e-08; Fig. 4E; significant in both animals [Table S4]). These effects were independent of the expected value of the current offer. Changing the history window between 3 and 10 trials did not change these conclusions.

We then asked whether focal serotonin delivery to the pgACC might influence the relationship between environment value and pursue-reject decisions. We observed that focal serotonin delivery to the pgACC significantly decreased the influence of environment richness on the likelihood of responding to a given reward opportunity (GLM2.2; β_MB*5-HT*Env.richness_ = −0.33, SE = 0.10, p = 0.00116; Fig. 4G). Effects were consistent across animals, with subject-level models showing the same direction of effect (Table S4). Thus, while animals normally adjusted their reward pursuit according to recent reward history, serotonergic manipulation of the pgACC attenuated this contextual sensitivity. Importantly, this effect was selective: focal serotonin delivery changed the effect of the general reward environment but not the effect of the expected value of each specific opportunity (GLM2.1; β_MB*5-HT*expected-value_ = 0.053, SE = 0.09, p = 0.550; Fig. 4H). These findings suggest that focal serotonin delivery to the pgACC specifically disrupts the influence of the statistics of the recent reward environment on ongoing decisions, without impairing the effect of immediate offers.

### A hidden Markov Model identifies latent motivation states

Behaviour exhibited significant autocorrelation across trials (Fig. 5A), suggesting the presence of state-like dynamics beyond immediate offer value. Accordingly, following previously outlined procedures for this task (17), we fitted a general linear model (GLM)-hidden Markov model (HMM) to identify latent motivation states in behaviour – defined as intrinsic variation in willingness to pursue reward that persists across trials, independent of the value of the current offer.

**Figure 5.**
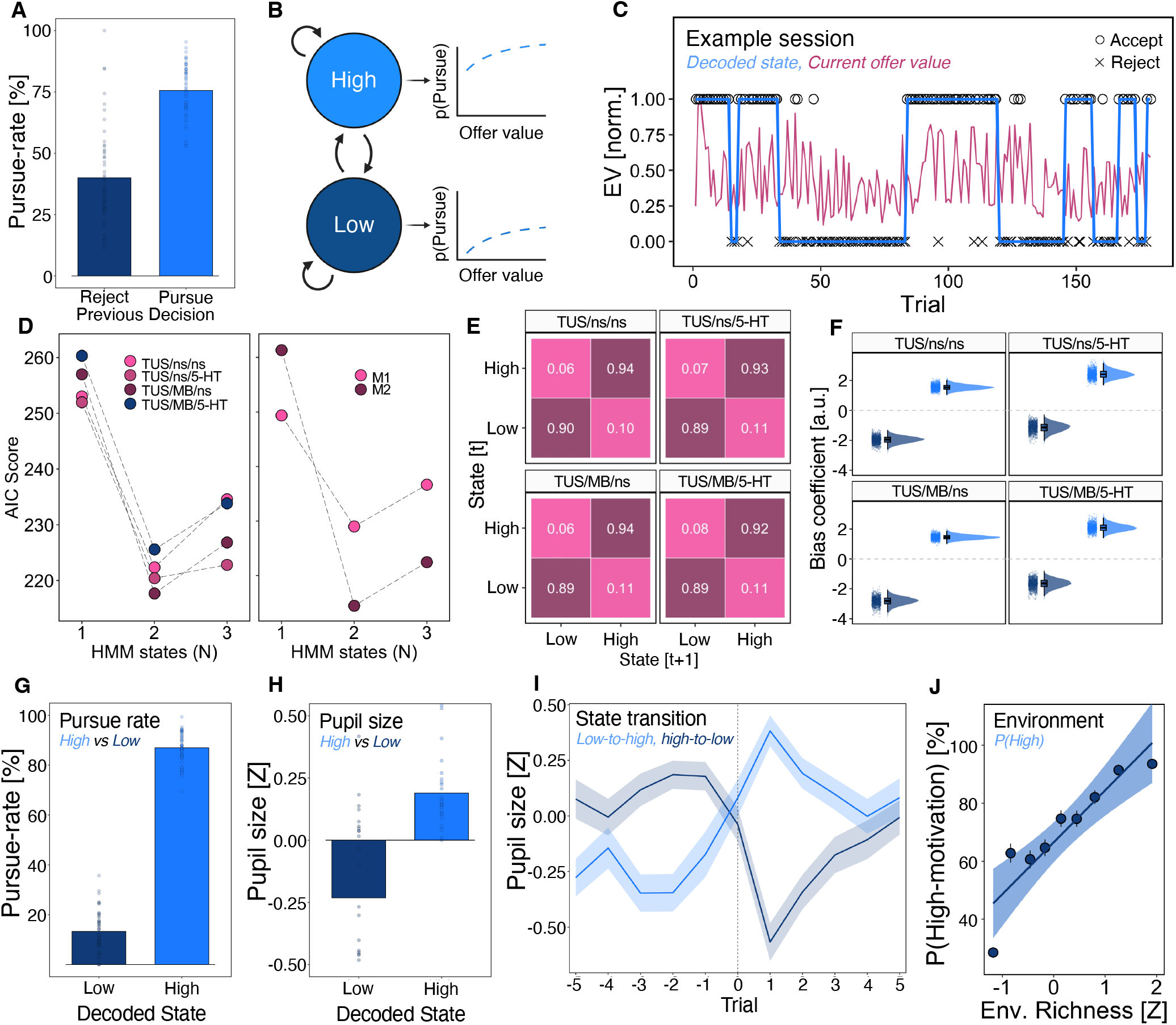
A GLM-HMM identifies latent motivation states. (A) Autocorrelation of pursue–reject behaviour across trials, indicating state-like temporal structure beyond immediate offer value. Each dot represents the mean pursue-rate for a single session, grouped by the previous trial’s decision. (B) Schematic of the GLM–HMM. Patterns in behaviour were captured with a GLM-HMM with state dependent bias parameters. These bias parameters produced state-dependent decision functions that reflected changes in the way animals made pursue-versus-reject decisions over time. (C) Example session illustrating decoded latent state trajectory (blue) overlaid on normalised expected value of the current offer (pink). Observed behaviour is shown for each trial, with pursue decisions above the trace and reject decisions below. (D) Model comparison showing that a two-state GLM–HMM provides a better fit relative to one and three state models according to AIC – a penalized form of log-likelihood – for each monkey and experimental condition. (E) State transition matrices for each condition, demonstrating high temporal persistence (autocorrelation) of inferred states across consecutive trials. (F) Posterior probability distributions of state-specific bias parameters for each condition. Half-violin plots show the distribution of bias coefficients across observations; boxplots indicate median and interquartile range, and points represent individual samples. Clear separation between states indicates meaningful differences in baseline propensity to pursue reward. (G) Animals were highly likely to pursue reward opportunities in a high motivation state, compared to low. Each dot represents the mean pursue-rate in a single session for high and low motivation states. (H) Pupil diameter as a function of decoded motivational state, showing significantly larger pupil size in the high- versus low-motivation state. Each dot represents the mean pupil size for each motivation state in a single session. (I) Trial-aligned pupil dynamics around decoded state transitions. Pupil size decreases prior to high-to-low transitions and increases prior to low-to-high transitions, with changes emerging 1–2 trials before the inferred transition. Shaded regions indicate ± SEM around the mean pupil size. (J) Probability of occupying the high-motivation state as a function of environment richness. Richer reward environments increase the likelihood of occupying the high-motivation state. Points indicate means of p(High) for deciles of environment richness; whiskers show normal-approximation 95% confidence intervals. Shaded region indicates the 95% confidence interval around the linear fit.

The GLM-HMM consists of a state-dependent GLM coupled to an HMM, incorporating predictors for the offer value, block cues, and an intercept (bias) term (Fig. 5B). To isolate latent motivational fluctuations, only the intercept term was allowed to vary across states, while other coefficients were constrained to be identical across states. Thus, latent states differed in baseline propensity to pursue offers while maintaining identical sensitivity to the value of the current offer.

An example session illustrates the trajectory of decoded states overlaid on trial-by-trial offer value and observed behaviour (Fig. 5C), demonstrating both the presence of latent state-like fluctuations and the model’s ability to capture these dynamics. Model comparison confirmed that a two-state GLM-HMM outperformed one- and three-state alternatives for each monkey and condition (Fig. 5D). Parameter recovery analyses further confirmed that model parameters could be reliably estimated (Fig. S3). This modelling framework has been validated in a closely related task (17), and here we use it to test how focal serotonergic manipulation of the pgACC alters motivational state dynamics.

To investigate the effect of focal serotonin delivery, separate GLM-HMMs were fit for each condition in the 2×2 design (Fig. 1D). Data from both monkeys were pooled within each condition, allowing the model to identify latent states that capture shared structure across animals. Each model returned a transition matrix (Fig. 5E), and state-specific bias terms (Fig. 5F). The bias terms show the models’ separation of high and low motivation states. Latent states were decoded using the Viterbi algorithm to obtain the maximum a posteriori state assignment on each trial (see Methods).

The high-motivation state was characterized by an increased likelihood of pursuing offers beyond what could be explained by current offer value (GLM3.1; β_state_ = 3.77, SE = 0.06, P < 2e-16; Fig. 5G; significant in both animals [Table S4]). Pupil diameter was also significantly larger in the high-motivation state than in the low-motivation state (GLM3.2; β_state_ = 408.02, SE = 28.52, p < 2e-16; Fig. 5H), providing an independent physiological readout of the inferred latent states.

Interestingly, pupil dynamics also anticipated state transitions. Pupil size began to change prior to decoded transitions between motivational states (Fig. 5I). Specifically, pupil diameter decreased preceding transitions from high to low motivation states and increased preceding transitions from low to high motivation states. These predictive changes were observable during the inter-trial interval, such that upcoming state transitions could be inferred from the interaction effect between the previous trial’s pupil size and state (GLM3.3; β_state*pupil-lag_ = −0.688, SE = 0.119, p = 7.97e-9). These findings demonstrate that the identified motivational states correspond to biologically meaningful, directionally consistent changes in pupil size that precede and predict behavioural state shifts.

We next asked what drives transitions between motivational states. If these transitions reflect animals’ reward context, recent reward should bias state occupancy. Indeed, the environment richness significantly increased the probability of occupying the high-motivation state (GLM3.4; β_Env.richness_ = 1.135, SE = 0.0298, p < 2e-16; Fig. 5J; significant in both animals [Table S4]).

These results identify latent, biologically grounded motivation states that integrate recent reward context beyond current offer value. This modelling approach provides an effective framework for testing how focal serotonin delivery to the pgACC shapes state transitions and motivated behaviour.

### Focal serotonin delivery to the pgACC disrupts context-dependent motivational state dynamics

Latent motivation state strongly influenced the likelihood of responding to a given offer. Under baseline conditions, richer reward environments increased the probability of occupying a high-motivation state, which in turn increased the likelihood of pursuing reward opportunities. We next examined how focal serotonin delivery to the pgACC altered these relationships.

Mixed-effects GLMs were used to quantify the effects of serotonergic manipulation on state occupancy and its dependence on reward context. Models included a by-monkey random intercept and effects were confirmed in subject-level analyses (Table S4).

Focal serotonin delivery to the pgACC significantly reduced the likelihood of occupying a high motivation state (GLM4.1; β_MB*5-HT_ = −0.26, SE = 0.087, p = 0.0026; Fig. 6A). In addition, serotonin delivery reduced the influence of environment richness on high-motivation state occupancy (GLM4.2; β_MB*5-HT*Env.richness_ = −0.775, SE = 0.122, p = 2.38e-10; Fig. 6B). Subject-level analyses confirmed that these effects remained significant when models were fit separately for each animal (Table S4).

**Figure 6.**
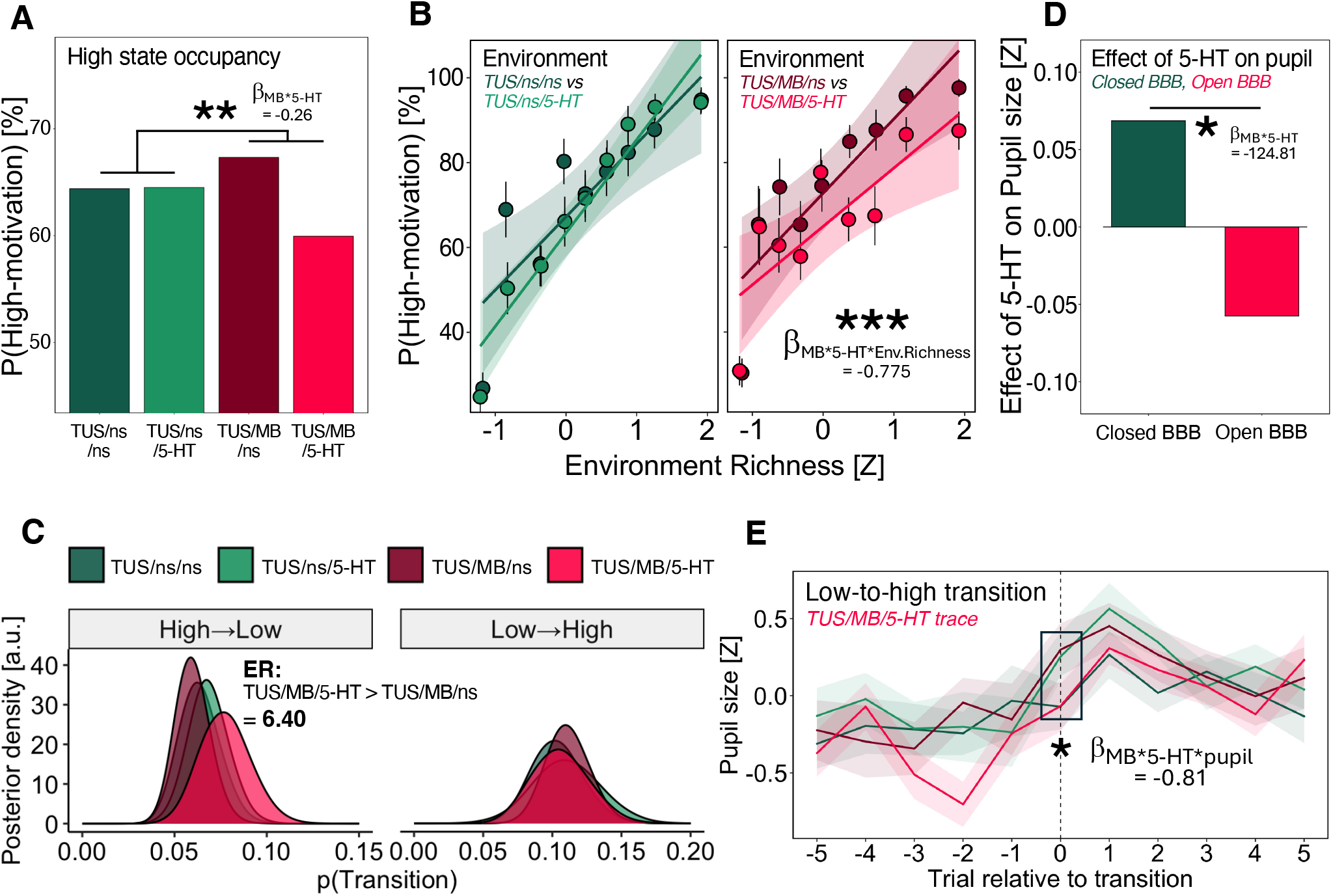
Focal serotonin delivery to the pgACC alters motivational state dynamics and associated pupil responses. (A) Probability of occupying the high-motivation state across conditions. Focal serotonin delivery (TUS/MB/5-HT) significantly reduced high-state occupancy relative to BBB opening alone (GLM4.1; β_MB*5-HT_ = −0.26, SE = 0.087, p = 0.0026). (B) Influence of environment richness on state occupancy. TUS/MB/5-HT significantly attenuated the effect of environment richness on the likelihood of being in a high motivation state (GLM4.2; β_MB*5-HT*Env.richness_ = −0.775, SE = 0.122, p = 2.38e-10). Points indicate means of p(High) for deciles of environment richness; whiskers show normal-approximation 95% confidence intervals. (C) Posterior probability of state transitions in GLM-HMMs fitted to each condition. Focal serotonin delivery (TUS/MB/5-HT) increased high-to-low transition probability relative to BBB opening alone (TUS/MB/ns) (P(delta > 0) = 0.865, evidence ratio (ER) = 6.40). (D) Pupil diameter across conditions. Systemic serotonin alone (TUS/ns/5-HT vs. TUS/ns/ns) increased pupil size, whereas focal serotonin delivery (TUS/MB/5-HT vs. TUS/MB/ns) was associated with a decrease in pupil diameter. (E) Trial-aligned pupil dynamics around decoded state transitions, split by condition. Compared to BBB opening alone (TUS/MB/ns), focal serotonin delivery (TUS/MB/5-HT) attenuated the predictive relationship between pupil size and upcoming low-to-high transitions (GLM4.4; β_MB*5-HT*pupil_ = −0.93, SE = 0.38, p = 0.0163).

To determine the mechanism underlying this shift in state occupancy, we examined transition dynamics between motivational states. Focal serotonin delivery to the pgACC increased the likelihood of transitioning from high-to-low motivation state relative to BBB opening alone (P(delta > 0) = 0.865, evidence ratio = 6.40; Fig. 6C), consistent with a reliable increase in high-to-low transitions, while leaving low-to-high transitions unchanged (P(delta > 0) = 0.416, evidence ratio = 0.71; Fig. 6C). No difference was observed (in the absence of BBB opening) between TUS/ns/5-HT and TUS/ns/ns conditions for either low-to-high or high-to-low transitions (Fig. 6C). This suggests that the reduced occupancy of the high-motivation state arises from an increased likelihood of transitioning out of that state.

Pupil dynamics further reflected these changes. Compared to baseline (TUS/ns/ns), systemic serotonin alone (TUS/ns/5-HT) increased pupil diameter (Fig. 6D). In contrast, under BBB opening conditions, serotonin delivery (TUS/MB/5-HT) was associated with a reduction in pupil size relative to BBB opening alone (TUS/MB/ns). This effect was captured by a significant negative interaction between serotonin and BBB opening (GLM4.3; β_MB*5-HT_ = −124.22, SE = 54.92, p = 0.0237; Fig. 6D) and remained significant after accounting for motivation state, indicating a distinct central effect of focal serotonergic manipulation detectable in pupil dynamics. Consistent with the state-dependent pupil signatures described above (Fig. 5I), focal serotonin delivery also altered the dynamics of pupil signals preceding state transitions. Specifically, compared to BBB opening alone, the anticipatory pupil increase in the inter-trial interval preceding low-to-high transitions was significantly attenuated (GLM4.4; β_MB*5-HT*pupil_ = −0.93, SE = 0.38, p = 0.0163; Fig. 6E). These effects were significant in one animal and showed the same direction of effect in the other (Table S4).

These results suggest that focal serotonin delivery at the pgACC disrupts the influence of the environmental value on motivation-state dynamics. By increasing transitions into low-motivation states and weakening the coupling between reward context and motivation state, serotonergic manipulation biases behaviour away from context-appropriate reward pursuit. These effects are accompanied by corresponding changes in pupil dynamics, providing an independent physiological confirmation of central serotonergic action rather than peripheral effects of systemic 5-HT.

### Repeated ultrasound-mediated neuromodulator delivery is well tolerated and increases serotonergic signal in the pgACC

Experiments 1 and 2 demonstrated both neural and behavioural effects of focal serotonin delivery. A central issue for BBB opening approaches, however, is whether repeated interventions induce cumulative tissue damage or sterile inflammatory responses that could limit translational potential (32,33). In Experiment 3, we used histology to assess tissue integrity following repeated BBB opening performed during Experiment 2, and to explore evidence for local delivery of serotonin at the target site.

Experiment 2 consisted of seven fortnightly blocks, each including two BBB opening sessions separated by at least 6 days. This resulted in 14 BBB opening procedures over approximately 4 months for each animal. Experiment 3 was conducted following the completion of Experiment 2 using the same animals (*n=2*), along with one control animal (*n=1*). Experiment 3 followed the same procedure as Experiments 1 and 2 but was performed under terminal general anaesthesia. BBB opening was followed by infusion of either serotonin (experimental group) or saline (control group).

No evidence of overt tissue damage, cellular loss, or lesioning was observed in either cresyl violet (CV) or haematoxylin and eosin (H&E) staining in animals that had undergone repeated BBB opening procedures (Fig. 7A,B).

**Figure 7.**
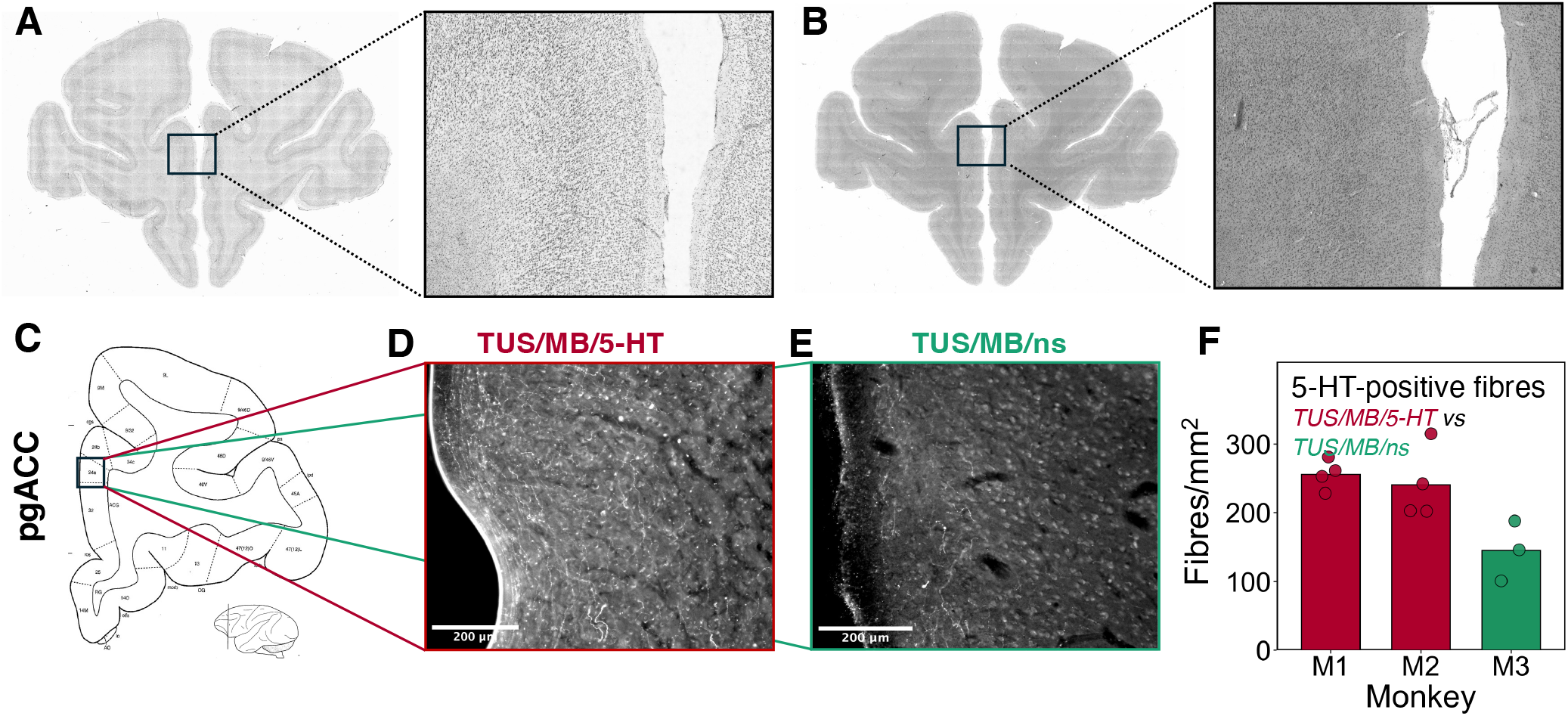
Focal serotonin delivery to the pgACC is well tolerated and increases serotonergic signal. (A,B) Cresyl violet (A) and haematoxylin and eosin (B) staining show no evidence of structural tissue damage in animals that underwent repeated BBB opening procedures at the pgACC. (C) Schematic of the histological sampling site corresponding to area 24a (34), a core component of the pgACC. (D,E) Representative sections showing serotonergic immunostaining in area 24a following TUS/MB/5-HT (D) and TUS/MB/ns (E). (F) Quantification of serotonergic labelling density (fibres/mm^2^) in area 24a. Animals receiving TUS/MB/5-HT (M1 & M2) show increased density of detectable serotonergic axonal fibres compared to BBB opening alone (M3). Each dot represents an individual histological section (M1, M2: 4 sections each; M3: 3 sections). Also see Fig. S4.

Finally, to provide direct evidence of local serotonin delivery, we performed immunostaining for serotonin in the pgACC (area 24a) following BBB opening (Fig. 7C-E; see methods) and in a control region without BBB opening (TUS without MBs; Fig. S4). Quantification revealed an increased density of serotonergic axonal labelling in pgACC of animals receiving focal serotonin delivery (TUS/MB/5-HT) compared to BBB opening alone (TUS/MB/ns; Fig. 7F). A comparable increase was not observed in the control region (Fig. S4), indicating spatial specificity of the effect. The increased serotonergic signal is consistent with enhanced local serotonin availability at the target site. Given that serotonergic axons express the serotonin transporter (SERT), this increase in immunoreactivity is likely to reflect uptake of delivered serotonin into the serotonergic fibres.

## Discussion

In this study, we demonstrated the use of ultrasound-mediated blood-brain barrier opening to deliver neuromodulators to a specific brain target in macaques. Using this approach, we delivered serotonin selectively to pgACC, altered local neural activity and connectivity, and showed that increasing serotonin at this site weakens the influence of environmental context on motivational state and behaviour.

We targeted pgACC because it is well placed to mediate serotonergic regulation of context-dependent decision-making. The pgACC receives direct input from DRN, is enriched in serotonergic receptors, and has been implicated in integrating reward history, volatility, and internal motivational variables to guide behaviour (35–37). Our findings support the idea that pgACC is a cortical site through which serotonergic signalling helps map environmental statistics onto adaptive behaviour.

Behaviourally, focal serotonin delivery to pgACC reduced the influence of reward environment on reward pursuit. In our task, richer environments normally increased the likelihood of occupying a high-motivation state, thereby increasing the probability of pursuing reward opportunities. Elevating serotonin at pgACC disrupted this relationship. Specifically, focal serotonin delivery reduced the influence of reward history on pursue-reject decisions, weakened the effect of recent reward context on high-motivation state occupancy, reduced the overall proportion of time spent in the high-motivation state, and increased the likelihood of transitions from high to low motivation states. Together, these results indicate that increasing serotonin in pgACC weakens the mapping from environmental statistics to latent motivational state and, in turn, to behaviour.

This effect was selective. We did not observe a change in sensitivity to the expected value of the current offer, nor did we detect an increase in low-to-high state transitions. Instead, serotonergic manipulation selectively perturbed the high-motivation state. The reduced occupancy of the high-motivation state arose because animals were more likely to transition out of it. This pattern suggests that serotonin at the pgACC does not broadly impair valuation but rather alters how recent environmental context is translated into sustained motivational engagement.

Pupil dynamics provided an independent physiological readout of these effects. Pupil diameter was larger in the high than in the low motivation state, and changed predictively before state transitions, increasing before low-to-high transitions and decreasing before high-to-low transitions. These dynamics indicate that the inferred motivational states are biologically meaningful. Focal serotonin delivery altered these physiological signatures: mean pupil diameter decreased, and the predictive pupil dynamics preceding state transitions were attenuated. These effects provide independent physiological confirmation that the behavioural changes arose from central serotonergic action rather than from nonspecific peripheral effects of systemic serotonin.

The neural data provided a candidate mechanism for this behavioural decoupling. Focal serotonin delivery reduced local glutamatergic metabolite signal in the pgACC and attenuated its resting-state functional connectivity with anatomically connected regions. This directional convergence suggests a shift toward reduced local excitatory tone and diminished engagement of the pgACC within broader frontal networks. Notably, BBB opening alone (without serotonin delivery) produced a small increase in functional connectivity, potentially driven by vascular or oxygenation changes post BBB opening, and although no effects were seen on MRS, implying limited metabolic effects in the region, this was accompanied by an upward trend in the proportion of time spent in the high-motivation state (31).

Serotonin immunostaining provided direct evidence of target engagement, revealing increased serotonergic labelling in the pgACC following focal serotonin delivery relative to BBB opening alone. This result is consistent with increased local serotonergic availability at the target site and supports the interpretation that the observed behavioural and neural effects arose from successful local delivery.

A plausible circuit-level account is suggested by the receptor architecture of pgACC. Serotonin acts through receptor- and cell-type-specific mechanisms in the cortex. In macaque anterior cingulate cortex, 5-HT_1A_ receptors are especially dense, with expression peaking in pgACC, whereas 5-HT_2A_ receptors are more broadly distributed (20,38). Because 5-HT_1A_ receptors are inhibitory and are concentrated in cortical input layers (39,40) – typically expressed on pyramidal glutamate neurons (41) – one possibility is that focal serotonin delivery suppresses activity in pgACC input circuitry, thereby reducing the impact of environmentally derived signals reaching this region. In parallel, serotonergic actions at 5-HT_2A_ receptors in deeper layers may reshape output signalling and synaptic integration (39,40), in line with the reconfiguration of pgACC functional connectivity reported above.

Importantly, although focal delivery likely elevated extracellular serotonin outside the normal physiological range, prior slice physiology shows that bath-applied serotonin produces robust, receptor-mediated changes in prefrontal/anterior cingulate pyramidal neurons, including 5-HT_1A_-mediated hyperpolarization and 5-HT_2_-mediated increases in excitability (42). This supports the interpretation that extracellular serotonin delivery can engage defined cortical serotonergic mechanisms. Given the high density of inhibitory 5-HT_1A_ receptors in the pgACC, the reduction in Glx signal and functional coupling observed here may reflect, at least in part, 5-HT_1A_-mediated suppression of local cortical excitability. We therefore propose that increased serotonin in the pgACC reduces the gain of contextual input while reshaping cortical output, producing the reduced glutamatergic signal, attenuated functional coupling, and weakened behavioural dependence on reward environment observed here.

Our findings also help position pgACC serotonin within a broader understanding of DRN. In related work (17), disruption of DRN reduced the influence of environmental value on motivational dynamics. The effect was similar in nature to the effect observed here – a reduction in the influence of the general reward environment rather than the current, specific opportunity. However, focal serotonin delivery to pgACC increased high-to-low transitions, whereas DRN disruption reduced these transitions. Notably, both manipulations altered high-to-low transitions while leaving low-to-high transitions relatively unchanged. These findings suggest that at least part of the DRN’s effect on motivational state is mediated through serotonergic projections to pgACC, and that this system may operate within an optimal range. Both decreasing and increasing serotonergic signalling may impair the effective coupling between environmental information and motivational state.

More broadly, these results support current accounts of serotonin as a regulator of transitions between behavioural states (17,43–46). Under normal conditions, serotonergic signalling from the DRN may enable cortical circuits to track changes in environmental value and maintain appropriate motivational engagement (47,48). In contrast, our manipulation likely elevated baseline serotonin levels in a spatially specific manner, reducing the dynamic range over which DRN signals can influence pgACC computations. As a result, elevated serotonin did not simply suppress behaviour, but weakened the mapping between environmental context, motivational state, and action.

Although our finding that elevating serotonin in the pgACC reduces occupancy of high-motivation states by weakening the influence of rich reward environments may appear at odds with the antidepressant efficacy of selective serotonin reuptake inhibitors (SSRIs), the two manipulations differ fundamentally. Here, we produced an acute, focal increase in serotonin within pgACC, whereas SSRIs induce chronic, brain-wide changes whose clinical effects depend on delayed neuroadaptive processes rather than serotonin elevation alone (49). Our results may therefore capture a circuit-specific component of serotonergic pharmacology that is also seen clinically, namely emotional blunting or reduced motivation in some patients receiving SSRIs (50). In this sense, serotonergic regulation of motivational state may be both circuit- and timescale-dependent, with both too little and too much serotonergic influence weakening the coupling between environmental value and behaviour.

Finally, these findings have important methodological implications. Repeated ultrasound-mediated BBB opening was well tolerated over months, with no evidence of tissue damage following multiple procedures, addressing key concerns for longitudinal applications. More broadly, this study establishes ultrasound-mediated neuromodulator delivery as a generalizable experimental platform for causal neuropharmacology in awake primates, expanding the range of questions that can be addressed in systems neuroscience and providing a foundation for future circuit-specific therapeutic interventions.

## Materials and Methods

### Structure and Timeline

We conducted three experiments. In Experiment 1, BBB opening and subsequent focal serotonin delivery were conducted in awake resting monkeys, without a behavioural task. In Experiment 2, BBB opening and focal serotonin delivery in awake monkeys were followed by performance of a behavioural task in the MRI scanner. Experiment 3 was conducted upon conclusion of Experiment 2, with focal serotonin delivery performed as part of a terminal procedure under anaesthesia after which histological analysis was conducted to assess safety and direct serotonin delivery.

In Experiments 1&2, we used a 2 × 2 factorial design manipulating MB administration and 5-HT infusion, with TUS applied in all conditions (Table S1). This design dissociated the effects of focal serotonin delivery to pgACC from those of serotonin infusion or BBB opening alone. Sessions were conducted in fortnightly blocks. No BBB opening session (TUS/MB/ns or TUS/MB/5-HT) followed another, ensuring a minimum interval of at least 6 days between BBB opening conditions to allow complete closure of BBB. Individual sessions were separated by at least 72 hours. Experiment 3 had two groups: BBB opening alone (TUS/MB/ns, control group) and BBB opening with serotonin infusion (TUS/MB/5-HT experimental group).

### Animals

Experiment 1 was conducted in three male rhesus macaques (*M. mulatta*), with one session per condition (*n=3* animals). Experiment 2 was conducted with two male rhesus macaques (*M. mulatta*), with seven sessions per condition (*n=2* animals). The animals were 14-16 years old and weighed between 15 to 17.5 kg. They were housed in groups on a 12-hour light-dark cycle. Access to water was *ad libitum* on non-testing days, and for 12-16 hours on testing days. Experiment 3 used the same macaques as in Experiment 2 (*n=2*), along with a control animal (*n=1*).

All procedures were conducted under a license issued by the UK Home Office in accordance with the Animal (Scientific Procedures) Act 1986 and the European Union guidelines (EU Directive 2010/63/EU).

### BBB opening and Experimental Procedure

Animals were positioned in an MRI-compatible chair in the sphinx posture. Intravenous access was obtained with a 22-gauge cannula placed in the saphenous vein, alternating legs when possible across the testing sessions. Animals were habituated to cannulation procedure to allow voluntary cannulation and intravenous infusion with minimal restraint.

Before TUS, the scalp was shaved and conductive gel applied (SignaGel; Parker Laboratories, Fairfield, NJ, USA). Acoustic coupling between the transducer and scalp was achieved using a water-filled coupling cone sealed with a latex membrane and filled with degassed water. The water was degassed for 4-6 hours before each stimulation session.

TUS was delivered using a 4-element annular array transducer (NeuroFUS CTX-500, 64mm active diameter, Brainbox Ltd., Cardiff, UK) driven by programmable amplifier (Sonic Concepts TPO-105, Brainbox Ltd., Cardiff, UK). TUS was applied with 500 kHz centre frequency, 10 ms pulse length and a 500 ms period. This corresponds to a 2 Hz pulse repetition frequency (PRF) and a 2% duty cycle. Stimulation proceeded for a total duration of 120s. Acoustic simulations were carried out in k-wave (51) to achieve a peak negative pressure of 0.4 MPa at the target. Stimulation parameters were chosen based on previously established protocols for safe and transient BBB opening in macaques (13). Targeting of the pgACC was performed using a neuronavigation system based on each monkey’s magnetic resonance imaging (MRI) scan of brain structure (Brainsight; Rogue Research, Montreal, QC, Canada).

For BBB opening, TUS was combined with intravenously administered perflutren lipid microbubbles (Luminity; maximum 6.4 × 10^9^ microspheres/ml). Microbubbles were activated immediately before use by shaking for 40 s (3M CapMix Capsule Mixing Unit). After venting vial pressure with a sterile needle, 0.3 mL (20µL/kg) microbubbles were mixed with 0.7 mL saline and administered starting 3 s before TUS onset over 20–25 s, followed by a 10 mL saline flush. In TUS/ns/ns and TUS/ns/5-HT conditions, the microbubble injection was replaced with a 10 mL saline bolus. To assess BBB opening at the pgACC, irrespective of the experimental condition, a gadolinium-based contrast agent (0.2 mL/kg Gadovist; gadobutrol 1.0 mol/L; Bayer) was injected immediately after TUS, followed by a 10 mL saline flush.

Following TUS, monkeys were moved into an MRI scanner and were connected to an infusion pump. Serotonin (Enzo serotonin hydrochloride) was infused intravenously at a target rate of 600 ng/kg/min for the whole duration of the scan (52). A 10 mM serotonin stock was prepared, and stored at −18 °C. This was diluted with saline to the working concentration determined by body weight and pump rate (0.2 mL/min). Body weights were monitored regularly to ensure accurate dosing. Infusion syringes were covered to protect serotonin from light. In TUS/ns/ns and TUS/MB/ns conditions, serotonin was replaced by normal saline and infused at the same rate.

All MR images were acquired with a horizontal bore clinical 3T scanner (Siemens MAGNETOM Prisma, Erlangen, Germany) with a 15-channel nonhuman primate–specific receive coil (RAPID Biomedical, Rimpar, Germany). Structural scans were acquired in both experiments and for every session to assess the leakage of the contrast agent into the brain. In Experiment 1, structural scan was followed by 1H-MRS and resting-state fMRI. In Experiment 2, structural scan was followed by task-based fMRI. The fMRI data from Experiment 2 is not reported in this manuscript. Total MRI scan time was typically 80 min. Experiment 3 procedure is detailed below.

At the end of the scans, the infusion was stopped, cannula was removed, and animals were returned to their home cage. There were at least 72 hours between each testing session. Conditions with BBB opening (TUS/MB/ns and TUS/MB/5-HT) did not follow each other, allowing sufficient time for BBB closure (14).

### Ultrasound Simulations

Ultrasound wave propagation for the TUS parameters was simulated following established procedures (53). Simulations were performed using k-Wave (51) to estimate peak pressure, spatial peak pulse-average intensity (ISPPA), and steady-state spatial pressure distributions. To achieve a peak negative pressure of 0.4 MPa at the pgACC, corresponding to an estimated focal intensity of 5.3 W/cm^2^, subject specific simulations indicated that a free-field ISPPA of 25 W/cm^2^ was required for M1, 30 W/cm^2^ for M2, and 25 W/cm^2^ for monkey M3.

Skull models were derived from pseudo-CT volumes generated from Black Bone MRI (26). The skull was segmented by thresholding between 1,400–2,100 Hounsfield Units (HU). Material properties (sound speed, density, absorption) were assigned by linear mapping from HU as described previously (54,55).

### Structural scan acquisition and processing

Contrast-enhanced T1-weighted structural images were acquired using an MP2RAGE sequence 10 minutes after the injection of the contrast agent (0.2 mL/kg Gadovist; gadobutrol 1.0 mol/L; Bayer). Imaging parameters were: isotropic 1.0 mm voxels (192 × 192 × 60 matrix; FoV = 192 mm); repetition time (TR) = 3000 ms; echo time (TE) = 3.59 ms; first inversion time (TI 1) = 500 ms; second inversion time (TI 2) = 2500 ms, echo spacing 9 ms; slice per slab = 60, phase-encoding = Right to Left.

To assess BBB opening, MP2RAGE data were reconstructed into INV1, INV2, and fitted T1 map (UNI images). INV1 images were used for quantification as they preserve greater sensitivity to contrast agent–induced T1 signal changes compared to the UNI image. INV1 images were brain extracted, bias corrected and registered to a common subject-specific space. BBB opening was confirmed by computing voxel-wise signal intensity ratios between sessions with BBB opening and matched sessions acquired within the same experimental round without BBB opening (TUS/MB/ns relative to TUS/ns/ns; TUS/MB/5-HT relative to TUS/ns/5-HT), within a subject-specific pgACC anatomical mask. Voxels exhibiting a signal increase greater than 20% were classified as demonstrating BBB opening (Table S2)

### Resting-state fMRI acquisition and preprocessing

rs-fMRI was collected in Experiment 1. Structural T1-weighted images from a previous dataset were used for registration (56). Functional images were collected with a CMMR multi-band gradient-echo EPI sequence with the following parameters: TR = 1282 ms; TE = 25.40 ms; flip angle = 63°; voxel size = 1.25 × 1.25 x 1.25 mm^3^; 42 interleaved coronal slices; FoV = 120 x 120 mm; multi-band acceleration factor = 2; GRAPPA acceleration factor = 2; phase-encoding direction foot-to-head (57). Each run comprised 1000 volumes (~21 min). Each animal completed each experimental condition once. One animal was excluded after preprocessing due to insufficient usable data, yielding 2 animals × 4 scans (one scan per condition) in the final analysis.

Preprocessing was performed using tools from FMRIB Software Library (FSL) (58), Advanced Normalization Tools (ANTs; http://stnava.github.io/ANTs) (59), Human Connectome Project Workbench (https://www.humanconnectome.org/software/connectome-workbench) (60), and the Magnetic Resonance Comparative Anatomy Toolbox (MrCat; https://github.com/neuroecology/MrCat). Preprocessing was done following procedures described previously (61).

Steps were:

1. Reorientation to radiological convention and removal of the first 10 s to reach steady-state RF excitation.
2. Motion correction with MCFLIRT using an average low-noise EPI as reference.
3. Outlier detection via symmetric mean absolute percentage error (sMAPE) between each volume and a reference EPI; thresholds were the minimum of a fixed cut-off and a distribution-based cut-off. Volumes exceeding threshold were flagged; runs of ≤2 consecutive flagged volumes were linearly interpolated. The longest continuous segment without unfixable outliers was retained; scans with <50% usable volumes after trimming were excluded.
4. Brain extraction, bias-field correction, and linear registration to the subject’s T1-weighted image.
5. High-pass filtering to remove scanner drift.
6. Extraction of white-matter and CSF time series; PCA to identify high-variance (noise) components, which were regressed out.
7. Low-pass filtering.

### Resting-state fMRI analysis

For seed-based analyses, preprocessed EPI time series were projected to the cortical surface and registered to the F99 macaque template using HCP Workbench (62,63). The medial wall was masked, data were smoothed with a 3 mm FWHM Gaussian kernel, the medial wall was re-masked, and time series were demeaned. Within each condition, data from individual animals were combined using group-PCA following procedures outlined in (64), retaining 200 principal components. This approach enhances signal-to-noise and captures variance components shared across animals, enabling estimation of condition-specific functional dynamics. Given the small number of subjects, these estimates reflect fixed-effects patterns and are not intended for population-level inference.

Regions of interest (ROIs) were defined based on known anatomical projection targets of the pgACC (29,30). Coordinates were obtained from the NIMH Macaque Template (NMT v2.0) and the Cortical Hierarchy Atlas of the Rhesus Macaque (CHARM) (65,66), aligned to MNI space, and transformed to F99 surface (67,68). A full list of ROI coordinates is provided in Table S3. Each ROI was defined as a 4 mm-radius circle on the cortical surface in both hemispheres and combined to form bilateral ROIs. Area V4 (dorsal, V4d) was chosen as a control.

For each seed, Pearson correlation coefficients were computed between the bilateral ROI-averaged time series and every cortical vertex. Correlations coefficients were Fisher z–transformed (clipped to [−2, 2]) and averaged across hemispheres. The resulting map represents average bilateral functional coupling between the seed ROI and the cortical surface.

Mean seed-to-target coupling was extracted for each predefined target ROI. These values were visualised as a fingerprint plot in which each angular step corresponds to a target ROI and the radial axis shows coupling strength.

To quantify global strengthening or weakening of connectivity across conditions, an ROI-wise difference (Condition 1 – Condition 2) in coupling strength was computed and summed across ROIs to yield a single ‘Net Connectivity Difference’ summary metric. This was used to compare the effect of 5-HT with the BBB open vs closed (i.e. between TUS/MB/5-HT – TUS/MB/ns vs TUS/ns/5-HT – TUS/ns/ns). Positive values indicated stronger average coupling in Condition 1; negative values indicated stronger coupling in Condition 2.

Statistical significance was assessed with a non-parametric permutation test. Subject-level label shuffling was infeasible with two animals. Condition labels were permuted independently within each ROI (without mixing data between ROIs), resulting in 2^12^ = 4096 permutations. P-values were corrected for multiple comparisons using the Benjamini–Hochberg false discovery rate method (α = 0.05).

### Magnetic resonance spectroscopy acquisition

1H-MRS was acquired in awake, head-fixed macaques (n=3). Spectra were localised to the perigenual anterior cingulate cortex (pgACC) using a semi-LASER sequence (CMRR Spectroscopy Package, University of Minnesota (69,70); TE = 60 ms, TR = 3000 ms, bandwidth = 1250 Hz, 1024 complex points, 512 transients, total acquisition time = 17:53 minutes). A cubic voxel (11 × 11 × 11 mm^3^; single oblique rotation of approx. 45°) was positioned using MP2RAGE structural images acquired earlier in the session to ensure accurate placement within cortical grey matter and to avoid recording from non-brain regions. Partial water suppression was done using VAPOR (71) (8 saturation pulses) with outer-volume suppression bands to reduce lipid contamination. The residual water signal was used for the alignment of transients before averaging them. An unsuppressed water reference was acquired from the same voxel location for eddy-current correction and as a reference input for spectral fitting. Raw data were exported in DICOM (.IMA) format and converted to NIfTI-MRS for processing with spec2nii from the FSL-MRS toolbox (72,73). Individual transients were retained for preprocessing. Individual coils were combined on the scanner using the supplied sequence-specific reconstruction pipeline.

### Magnetic resonance spectroscopy preprocessing and quantification

Preprocessing and quantification were performed with the FSL-MRS toolbox (version 2.4.6), following recommended single-voxel preprocessing guidelines (74) with prespecified adaptations described below. Briefly, partially water-suppressed transients were cleaned using automated outlier detection, then frequency/phase aligned (spectral registration between 1.9–5.0 ppm), averaged across transients, and eddy-current corrected using the matched water reference. Residual water was removed using HLSVD and spectra were phase corrected. Water-reference spectra were phase corrected. For scans with residual frequency offset, an additional frequency shift was applied to align the NAA singlet peak to 2.01 ppm.

Spectra were quantified in FSL-MRS using a basis set simulated using FSL’s density matrix simulation tool *fsl_mrs_sim*. Basis set simulation used non-idealised pulses, a full description of the sequence timings, and was spatially resolved (40 points). The following metabolites were included in the basis set: Ala, Asc, Asp, GPC, PCh, Cr, PCr, GABA, Glc, Gln, Glu, GSH, Ins, Lac, NAA, NAAG, PE, Scyllo, Tau, as well as an empirically measured metabolite-nulled macromolecular basis (measured from a matched sequence in human cortex). Spectra were fitted using a polynomial baseline of order 2. Concentrations of glutamate and glutamine were summed to estimate Glx. Metabolite concentration estimates were expressed as ratios to total creatine (Glx/tCr and NAA/tCr), and fit uncertainty was summarized using Cramér-Rao lower bounds (CRLB). No tissue-fraction correction was applied. Spectral quality was assessed using FSL-MRS output metrics including SNR and linewidth defined as the fitted tCr full width at half maximum (tCr FWHM). A representative spectrum is shown in Figure 3A. QC metrics are reported for transparency (Fig. 3C).

### Pupil size acquisition and preprocessing

Pupil diameter and gaze (x, y) were recorded with an MR-compatible EyeLink 1000 (SR Research) at 250 Hz. Data were preprocessed and analysed in R following a standard pipeline adapted from (75): eye blinks were identified using the EyeLink system’s native detection algorithm, and artefacts were defined as consecutive samples with changes exceeding 50 arbitrary units. Subsequently, samples within symmetric 25-sample windows (0.1 s) surrounding detected artefacts or blinks were removed and linearly interpolated. The resulting time series were low-pass filtered at 4 Hz. These steps were applied independently to pupil size, horizontal gaze (x), and vertical gaze (y) signals, after which each time series was z-scored. To account for variance attributable to gaze position, pupil size was regressed on x- and y-gaze coordinates using linear regression, and subsequent analyses were conducted on the residuals. Pupil size was quantified within specific epochs relevant to decision-making around reward opportunities. Specifically, for each trial, mean pupil size was computed from the 0.5 s prior to each stimulus onset (inter-trial interval, ITI). This phase was chosen to make sure that pupil measurements were not confounded by the brightness of the stimulus. GLMs involving pupil size are detailed below in the relevant behaviour section.

### Behavioural training

Task specific training was conducted as described previously (17). This present study used the same task. Briefly, all monkeys were fitted with MRI-compatible cranial implants enabling head fixation during testing and training. Training occurred in MRI-compatible chairs in a sphinx posture within custom-built mock-scanner environments. Animals first performed a simplified task without a go-cue, action/outcome delays, or reward-environment changes. Go-cue and action/outcome delays were then introduced and gradually increased to durations suitable for fMRI. Reward-environment manipulations were then added after animals adapted to the final delay timings.

### Behavioural task

Monkeys performed a sequential decision task with visual cues of reward opportunities presented on a computer display. Each opportunity appeared as a coloured, dot-filled rectangle (8 × 26 cm). Reward magnitude (smoothie) was encoded by colour (red = 1 drop; green = 2; blue = 3), and reward probability was encoded by number of dots (each dot = 0.05 probability increment). Opportunities initially appeared at the centre of the screen, then shifted to the left or right, with the displacement serving as the go cue. Animals pursued an opportunity by breaking an infrared sensor with their left or right hand on the corresponding side. Lateralisation was randomised to prevent motor planning during deliberation. The go-cue temporally separated choice from movement initiation. The interval from opportunity onset to go cue was drawn from a uniform distribution, U(3,4) s. Animals then had 1 s to respond. After a pursue response, action-outcome delays were drawn from U(3, 4) s before potential reward delivery and on-screen feedback. The inter-trial interval followed ITI ~ U(3, 4) s. Skipped opportunities advanced immediately to the next trial (no action-outcome or reward). Premature responses (before the go cue) incurred the remaining offer presentation time plus the full go-cue, action-outcome, and feedback intervals; the same opportunity was then repeated on the next trial.

Each session comprised four blocks of 40–50 trials. Blocks manipulated the reward-probability landscape along two axes:

- Mean probability of reward: μ_rich = 0.80 vs μ_poor = 0.55.
- Stochasticity of offers: predictable blocks drew probabilities from a Gaussian with σ_predictable = 0.05; unpredictable blocks drew from a random-uniform range of 0.40 (yielding σ_stochastic = 0.13).

These factors yielded four environments: rich–predictable, rich–stochastic, poor–predictable, and poor–stochastic. Each session included one block of each type. Block order was counterbalanced with respect to richness to avoid extended runs of low-value environments.

The task was written in MATLAB (2019; MathWorks) using Psychophysics Toolbox v3 (76) and presented on an MRI-compatible 23″ BOLD screen (Cambridge Research Systems) ~30 cm from the animal. Smoothie rewards were a water–blackcurrant squash–banana mix, each drop was 1mL. Animals completed one session per day at the same time.

### Behavioural analysis

Analyses of behaviour first investigated the predictors of binomial pursuit/reject decisions. Pursuit (coded as ‘1’) versus rejection (coded as ‘0’) was modelled with mixed-effects binomial GLMs that included subject as a random effect, so inter-subject variability was accounted for in all analyses. Given the small number of subjects (*n* = 2), it was not statistically possible to estimate full random-effects structures for all predictors, as this would result in unstable parameter estimates. We therefore adopted a conservative approach, including random intercepts by subject and allowing random slopes only where supported by the data and model convergence. To ensure robustness, all key effects were additionally verified in subject-level analyses.

Following (17), environment richness was defined as the five-trial retrospective moving average of reward rate.

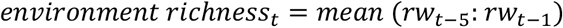

#### Exploratory GLMs (GLM1)

GLMs were run to explore the effect of predictors on pursue rate. These were: reward-magnitude (GLM1.1), reward-probability (GLM1.1), reward-expected-value (GLM1.2 & GLM1.3), and environment-richness (GLM1.3). Trial number (*trial*) was added as a confound regressor to control for the effect of fatigue. All mixed-effects GLMs were run in R and fit by maximum likelihood using lme4. GLMs 1.1-1.3 followed the same structure:

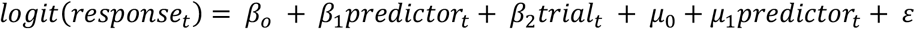

#### Characterising the effect of focal serotonin delivery on model-free behaviour (GLM2)

To probe the behavioural effects of serotonin (‘5-HT’; coded as 1 or 0) when opening the blood-brain barrier (‘MB’; coded as 1 or 0), these factors were included in GLMs 2.1 and 2.2, as in the 2×2 design (Fig. 1A).

GLM2.1 was run to investigate if focal serotonin delivery modulates the effect of the expected-value of the current offer on pursue rate. This was fit with fixed effects β_0_ + β_1_ expected-value* MB * 5-HT + β_2_ trial, a by-subject random intercept μ_0_ and an error term ε. In all GLMs including interaction terms, models were specified with the corresponding main effects and lower-order interactions.

***GLM2.1:***

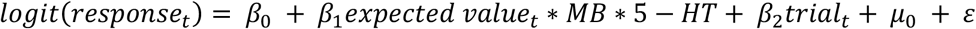

GLM2.2 investigated how focal serotonin delivery modulates the effect of environment richness on pursue rate, and was fit with fixed effects β_0_ + β_1_ expected-value + β_2_ environment-richness * MB * 5-HT + β_3_ trial, a by-subject random intercept μ_0_ and an error term ε.

***GLM2.2:***

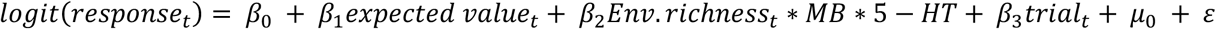

#### Modelling motivation states

As in (17), a generalized linear model-hidden Markov model (GLM-HMM) was used to capture slow, time-varying structure in behaviour consistent with internal motivation states. The HMM assumes that choices arise from two latent states, that evolve as a first-order Markov chain; the state (*z*) at trial *t* depends only on the state at *t−1* via a stationary transition matrix **A**, where *A_j,k_* is the likelihood of transitioning from state *j* to state *k*.

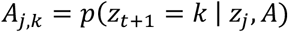

Conditional on the current state, choices follow a GLM whose parameters can be state-specific. On each trial *t*, the animal chose to pursue or reject the opportunity (binary outcome). To isolate motivational fluctuations, we allowed only the bias term to vary across states; the EV and environment-cue weights were constrained to be identical across states. Thus, states capture shifts in the baseline propensity to pursue, rather than changes in how evidence is weighted.

We fit models by Markov chain Monte Carlo in Stan. For each condition, trials from both monkeys were concatenated and a single GLM-HMM was fit to the combined dataset, allowing the model to identify latent states that capture shared structure across animals. To test the benefit of latent states, we compared K ∈ {1,2,3} and reported session-wise Akaike Information Criterion (AIC), which penalizes complexity and becomes stringent for K ≥ 2 as the number of parameters grows rapidly. To further validate the model, we tested whether parameters could be recovered from synthetic datasets generated from fitted 2-state GLM-HMMs for each condition. Generative parameters were set to the posterior means of the fitted models, and simulated datasets preserved the session structure and trial-wise covariates of the empirical data. For each session, we generated 10 simulated datasets. We then refit both 1- and 2-state models using the same Markov chain Monte Carlo procedure and compared fits using session-wise log-likelihood (computed via the forward algorithm) and AIC. We additionally assessed recovery of the transition matrix and state-specific bias terms by comparing recovered posterior distributions to the generative parameters. Subsequently, the maximum *a posteriori* sequence of latent HMM states in each session was decoded using the Viterbi algorithm, where *z_1:T_* denotes the sequence of latent states over time steps *1:T*, *y_1:T_* denotes the corresponding observed data, and *z\** denotes the most probable state sequence given the observations:

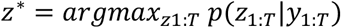

Each model output a transition matrix A, and the bias coefficients for each state. Transition trials were defined as the first trial of a new decoded state. The distribution of high-to-low and low-to-high transition probabilities for each model (and thus each condition) are visualised in Fig. 5E. To compare a parameter across models, we compute the posterior distribution of their difference and assess deviation from zero. Evidence ratios were used to quantify this deviation, defined as the ratio of posterior probability mass supporting positive versus negative values of the difference (i.e. ER = P(Δ>0)/P(Δ<0)), which provided a measure of directional evidence for parameter differences.

#### Characterising state-based behaviour (GLM3)

Further GLMs were run to characterise the effect of motivation-state on various behaviours of interest. The decoded state (High or Low) was included as a factor. GLM3.1 was run to investigate if motivation-state modulated the pursue rate. This was fit with fixed effects β_0_ + β_1_ expected-value + β_2_ state + β_3_ trial, by-subject random intercepts μ_0_ and μ_1_ state, and an error term ε.

***GLM3.1:***

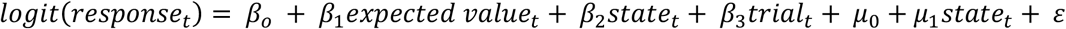

GLM3.2 investigated if pupil size was modulated by motivation-state. This was fit with fixed effects β_0_ + β_1_ expected-value + β_2_ state + β_3_ trial, a by-subject random intercept μ_0_, and an error term ε.

***GLM3.2:***

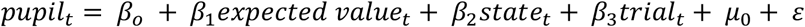

GLM3.3 was run to probe whether changes in pupil size could predict upcoming transitions. This was fit with fixed effects β_0_ + β_1_ pupil-size on the previous trial + β_2_ state + β_3_ trial + β_4_ monkey.

***GLM3.3:***

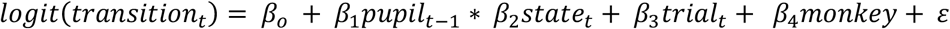

GLM3.4 investigated the effect of environment richness on being in a high motivation state. This was fit with fixed effects β_0_ + β_1_ expected-value + β_2_ environment-richness + β_3_ trial, a by-subject random intercept μ_0_ and an error term ε.

***GLM3.4:***

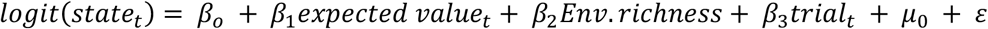

#### Characterising the effect of MB*5-HT on state-based behaviour (GLM4)

Changes in decoded HMM states were used to examine the temporal dynamics of motivation states. See the previous section for further information on Viterbi state decoding. We were interested in links between motivation states and aspects of the external environment, such as the recent reward history (environment richness).

GLM4.1 investigated how MB*5-HT changes the likelihood of being in a high motivation state, and was fit with fixed effects β_0_ + β_1_ expected-value + β_2_ MB * 5-HT + β_3_ trial, a by-subject random intercept μ_0_ and an error term ε. As above, all interaction terms included the corresponding main effects and lower-order interactions.

***GLM4.1:***

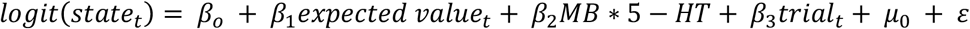

GLM4.2 investigated how MB*5-HT modulates the effect of environment on being in a high motivation state. GLM4.2 was fit with fixed effects β_0_ + β_1_ expected-value + β_2_ environment-richness * MB * 5-HT + β_3_ trial, a by-subject random intercept μ_0_ and an error term ε.

***GLM4.2:***

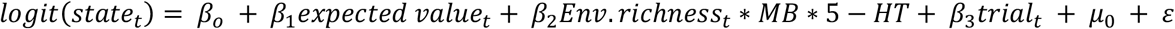

GLM4.3 investigated how MB*5-HT modulated pupil size. GLM4.3 was fit with fixed effects β_0_ + β_1_ state + β_2_ MB * 5-HT + β_3_ trial, a by-subject random intercept μ_0_ and an error term ε. ‘state’ was included in the model to ensure that any obtained effect from the MB*5-HT on pupil size is over and above potential effect of motivation states on pupil size.

***GLM4.3:***

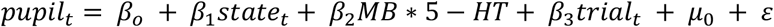

GLM4.4 investigated how MB*5-HT modulated the effect of pupil size on predicting low-to-high state transitions. GLM4.4 is fit with fixed effects β_0_ + β_1_ pupil * MB * 5-HT + β_2_ trial + β_3_ monkey and an error term ε.

***GLM4.4:***

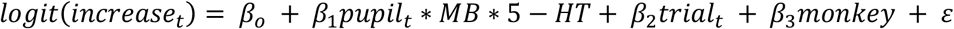

### Post-mortem Analyses (Experiment 3)

In Experiment 3, ultrasound-mediated BBB opening was conducted as described above, although under anaesthesia instead of awake, followed by either serotonin or saline infusion.

#### Animals

Brains of three male monkeys were collected post-mortem for histological analysis. Two monkeys (experimental group, n = 2) received focal serotonin delivery following BBB opening at pgACC (TUS/MB/5-HT) and TUS alone at a control region (visual cortex; TUS/ns/5-HT). The control monkey (control group, n = 1) received normal saline following BBB opening at pgACC (TUS/MB/ns) and TUS alone at a control region (visual cortex; TUS/ns/ns). Animals were sedated with triple combination of ketamine, medetomidine, and midazolam and maintained on sevoflurane low concentration 1 to 1.5% and ketamine infusion (5-10 mg/kg/h) to maintain immobility and limit the BBB perturbation.

#### Perfusion and tissue processing

After termination of the serotonin (or saline) infusion, animals were transcardially perfused with phosphate-buffered saline (PBS) until the effluent was clear, followed by 4% paraformaldehyde (PFA) in PBS. Brains were post-fixed in 4% PFA for 24 h at 4 °C and stored in PBS containing 0.02% sodium azide. Brains were sectioned coronally at 40 μm using a base sledge microtome (Leitz-Wetzlar; model 1300) and stored in cryoprotective solution. It took between 10-20 min between termination of the serotonin (or saline) infusion and the tissue fixation.

#### Cresyl Violet and H&E staining

Every fifth section within the pgACC (every tenth section for the control region), defined using the Paxinos atlas (34), was processed for cresyl violet (CV) or haematoxylin and eosin (H&E) staining. Sections were taken through graded alcohols, stained using standard CV or H&E protocols (including haematoxylin differentiation and bluing for H&E), dehydrated, cleared in Histoclear, and coverslipped with mounting medium.

#### Immunohistochemistry

Free-floating sections (every fifth section for pgACC; every tenth for control region) were washed in PBS and treated with 0.5% sodium borohydride, followed by incubation in glycine/PBST and blocking solution (PBST with 5% normal goat serum). Sections were incubated with rabbit anti-serotonin primary antibody (ImmunoStar, #20080; 1:5000) for 1 h at room temperature and overnight at 4 °C. After washing, sections were incubated with Alexa Fluor 568-conjugated goat anti-rabbit secondary antibody (Abcam, ab175471; 1:500) for 2 h. Nuclei were counterstained with DAPI (1:1000). All fluorescent steps were performed protected from light. Sections were mounted and coverslipped with fluorescence mounting medium (Abcam ab104135).

#### Imaging and Quantification

Brightfield (CV and H&E) sections were imaged using a Zeiss Axio Imager Z.1 microscope (5× and 20× objectives). Immunofluorescent sections were imaged using an epifluorescence microscope (Olympus BX40) with consistent acquisition settings across samples. Serotonergic fibres were quantified within the pgACC, corresponding to area 24a as defined by the Paxinos atlas (34), and the control region. Quantification was performed using ImageJ on multiple histological sections per animal (M1 and M2: 4 sections each; M3: 3 sections). For each section, regions of interest were manually defined within the pgACC and the visual cortex, and serotonergic axonal profiles were identified and counted. Axon counts were initially obtained separately for the left and right hemispheres. As no hemispheric differences were observed (paired *t*-test, *p* = 0.778), counts were combined across hemispheres for subsequent analysis. Axonal density was calculated by normalising axon counts to the sampled area (fibres/mm^2^).

### Analysis of hippocampal synaptic transmission following TUS

We investigated any potential neuromodulatory effects of TUS alone (without MB) using rodent hippocampal slice electrophysiology. Extracellular field recordings were performed in acute hippocampal slices from juvenile male and female Wistar rats. A recording electrode was placed in the *stratum radiatum* of the CA1 and a bipolar stimulating electrode was positioned in the Schaffer collateral. Synaptic responses were evoked by a single stimulus delivered at increasing intensities (0–20 V, 1 V increments; 0.067 Hz). At each intensity, two consecutive responses were recorded and averaged. Presynaptic fibre volley (FV) amplitude and the slope of the field excitatory postsynaptic potential (fEPSP) were measured. Input–output relationships were generated by plotting FV amplitude and fEPSP slope against stimulus intensity, and by plotting fEPSP slope as a function of FV amplitude to normalise for presynaptic recruitment. Data are presented as mean ± SEM across slices. Linear or nonlinear regression models were fitted as appropriate, and differences between conditions were assessed using Extra Sum of Squares F-tests.

## Supplementary Figures

**Supplementary Figure 1.**
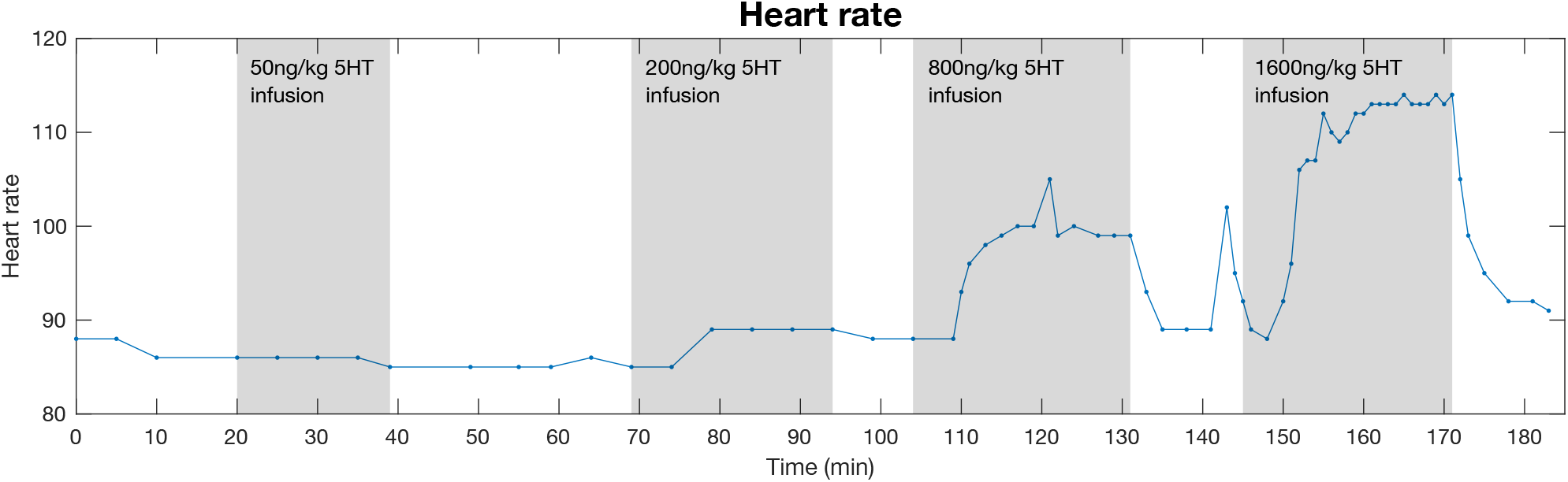
Dose determination experiment. Heart rate trace from a dose-determination experiment conducted under anaesthesia in one animal, with continuous veterinary monitoring of heart rate, blood pressure, and temperature. A range of serotonin (5-HT) infusion rates, selected based on prior work (52) was tested to identify a dose that produced measurable peripheral effects while remaining within a safe physiological range. A dose of 600 ng/kg/min was selected for subsequent experiments based on these data and previous studies (52).

**Supplementary Figure 2.**
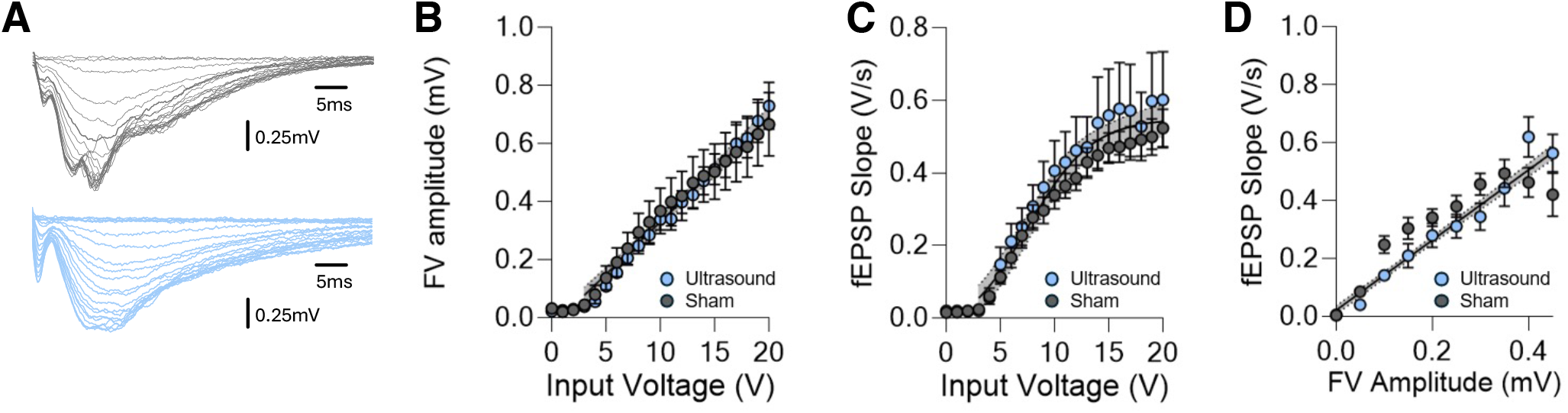
Effect of TUS parameters on ex vivo hippocampal synaptic circuit function. (A) Representative voltage traces from CA1 field recordings in response to incremental stimulation (0–20 V) of the SC pathway from sham (grey) and US (blue) treated slices. (B) Presynaptic fibre volley (FV) amplitude as a function of stimulation intensity. (C) Field EPSP (fEPSP) slope plotted against stimulation intensity. (D) fEPSP slope plotted against FV amplitude (0.05 mV bins) to normalise for presynaptic recruitment. Data represent mean [± SEM] across slices. Lines indicate regression fits (shaded area is 95% CI). Difference between model fits analysed by Extra Sum of Squares F-test finding no significant difference between ultrasound (blue) and sham (grey) conditions (B: sham: 0.34 ± 0.04 mV/ms μV^−1^ vs. ultrasound: 0.33 ± 0.1 mV/ms μV^−1^; DFd = 626, F = 0.5, p = 0.60; C: sham: 0.29 ± 0.05 mV/ms μV^−1^ vs. ultrasound: 0.34 ± 0.05 mV/ms μV^−1^; DFd = 622, F = 2, p = 0.69; D: sham: 0.32 ± 0.05 mV/ms μV^−1^ vs. ultrasound: 0.32 ± 0.06 mV/ms μV^−1^; DFd = 701, F = 2.75, p = 0.65). N = 8 rats; 15 slices per condition.

**Supplementary Figure 3.**
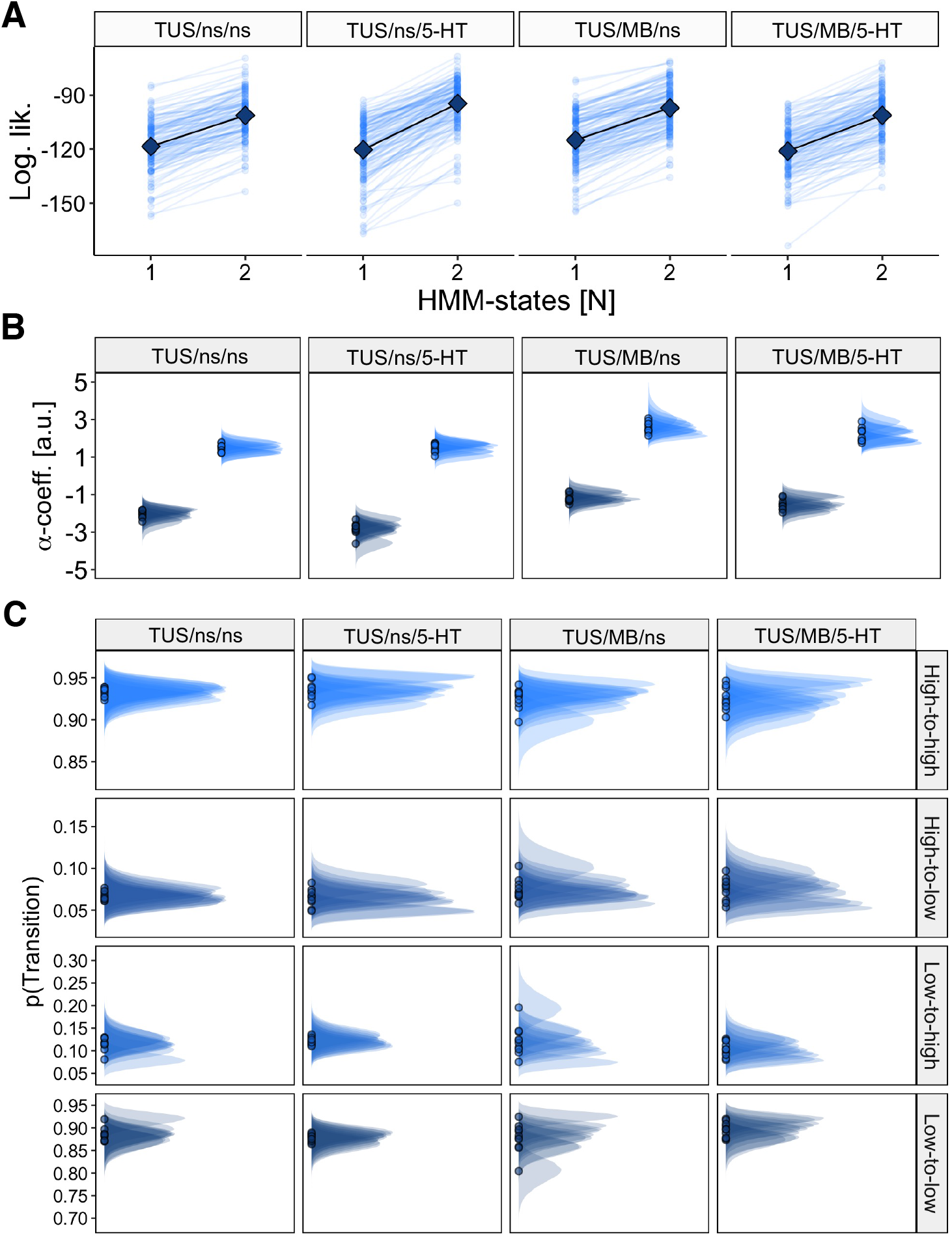
HMM-GLM Parameter Recovery. We assessed the performance of fitting 2-state GLM-HMMs by generating synthetic data from our models and subsequently refitting both 1 and 2 state models to this synthetic data. (A) Session-wise log-likelihoods for models with different numbers of latent states fitted to simulated data generated from a 2-state GLM-HMM. Thin blue lines show individual session fits across simulations; black points/lines indicate the mean across sessions. Data are shown separately for each experimental condition (TUS/ns/ns, TUS/ns/5-HT, TUS/MB/ns, TUS/MB/5-HT). Higher log-likelihood for the 2-state model demonstrates reliable recovery of the true latent state dimensionality across conditions. (B) Posterior distributions of state-specific intercepts (α) from the GLM-HMM fitted to simulated data. Each distribution corresponds to one latent state (low vs high) and condition. Clear separation between states and consistency across simulations indicate robust recovery of state-dependent choice biases. (C) Posterior distributions of transition probabilities between latent states recovered from simulated datasets. Distributions are shown for each transition type (low-to-low, low-to-high, high-to-low, high-to-high) and condition. Consistent clustering of samples indicates accurate recovery of the underlying Markov transition structure across simulations and experimental manipulations.

**Supplementary Figure 4.**
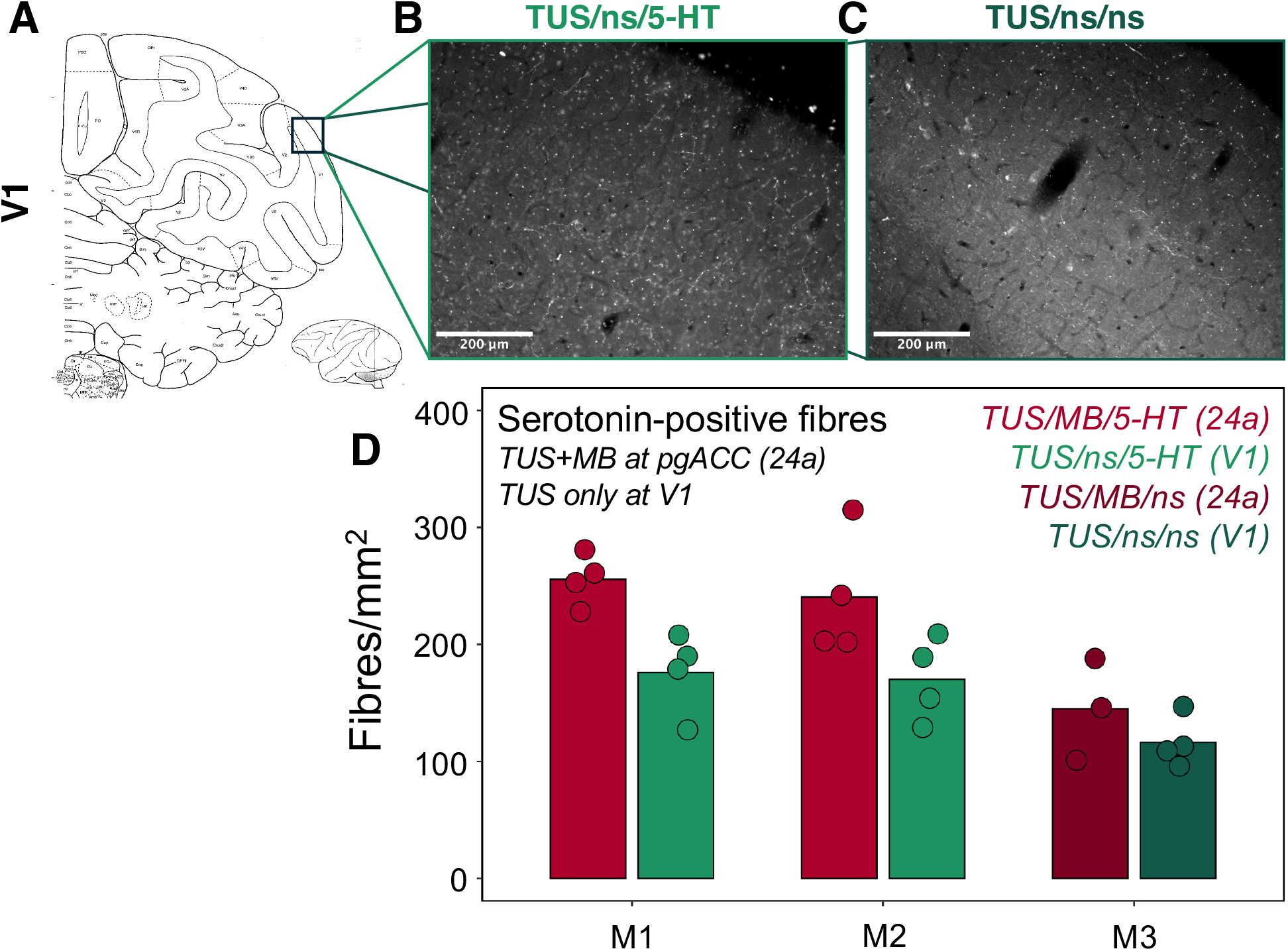
Quantification of serotonin-positive fibres in control region. (A) Schematic of the V1 control sampling site. (B,C) Representative sections showing serotonin immunostaining in V1 following TUS/ns/5-HT (B) and TUS/ns/ns (C). (D) Quantification of serotonin-positive fibres (fibres/mm^2^) in area 24a (pgACC) and primary visual cortex (V1), where TUS was applied without MBs. Animals receiving TUS/MB/5-HT at pgACC (M1, M2; experimental group) showed a clear increase in density of detectable serotonergic axonal fibres in area 24a relative to V1 (TUS/ns/5-HT). A comparable increase was not observed in the BBB opening–only condition (M3; control group; TUS/MB/ns vs. TUS/ns/ns). Each dot represents an individual histological section.

## Supplementary Tables

**Supplementary Table 1.** 2×2 Study Design.

| Condition | TUS | Microbubbles | Infusion |
| --- | --- | --- | --- |
| TUS/ns/ns | Yes | No | Normal saline |
| TUS/ns/5-HT | Yes | No | Serotonin |
| TUS/MB/ns | Yes | Yes | Normal saline |
| TUS/MB/5-HT | Yes | Yes | Serotonin |

**Supplementary Table 2.** Session-wise volume (mm^3^) of contrast-enhanced voxels after BBB opening. Volume was calculated by summing voxels with a 20% increase in signal from sessions with BBB opening and matched sessions acquired within the same experimental round without BBB opening (TUS/MB/ns relative to TUS/ns/ns; TUS/MB/5-HT relative to TUS/ns/5-HT), within a subject-specific pgACC anatomical mask. The mean BBB opening volume across sessions was 87 mm^3^. Sessions marked as N/A indicate cases in which contrast agent administration was not performed or was affected by issues related to dose or timing, precluding reliable quantification of contrast enhancement.

|  | M1 |  | M2 |  |
| --- | --- | --- | --- | --- |
| *experimental block* | TUS/MB/ns | TUS/MB/5-HT | TUS/MB/ns | TUS/MB/5-HT |
| rs-fMRI | 103 | 138 | 43 | 33 |
| 1 | N/A | 94 | N/A | 36 |
| 2 | 55 | 110 | 2 | 57 |
| 3 | 64 | 7 | 16 | 61 |
| 4 | 122 | 64 | N/A | 18 |
| 5 | 108 | N/A | 190 | N/A |
| 6 | 175 | 126 | 5 | 249 |
| 7 | 144 | 120 | 110 | 85 |

**Supplementary Table 3.** Regions of Interest (ROIs) used in resting-state fMRI analysis. Coordinates are in MNI macaque space.

| Area | Abbreviation | MNI coordinates [x,y,z] (67) |
| --- | --- | --- |
| Perigenual anterior cingulate cortex | pgACC | -1.5 17.5 7.5 |
| Subgenual anterior cingulate cortex | sgACC | -0.86 10.05 -1.27 |
| Mid-cingulate cortex | MCC | -0.92 -3.55 11.36 |
| Posterior cingulate cortex | PCC | -0.92 -14.47 9.54 |
| Anterior superior temporal gyrus | aSTG | -23.0 0.55 -6.14 |
| Middle superior temporal sulcus | mSTS | -16.8 -10.39 -5.09 |
| Anterior insula | AI | -16.44 3.3 -1.25 |
| Rostral entorhinal cortex | rERh | -6.92 -1.16 -14.16 |
| Area 9 medial | 9m | -1.5 19 12.5 |
| Area 9/46d dorsal | 9-46d | -12.75 11.5 9.25 |
| Area 11 medial | 11m | -4.14 21.98 1.89 |
| Area V4 dorsal | V4d (control) | -18.19 -26.50 12.49 |

**Supplementary Table 4.** Selected key GLMs run for individual monkeys. The predictor of interest is shown in bold. Each row shows beta estimates & p-values for that predictor for each monkey.

| Selected GLMs |  | M1 |  | M2 |  |
| --- | --- | --- | --- | --- | --- |
| | | $\beta$ | $p$ | $\beta$ | $p$ |
| 1.2 | *response ~ trial + **EV*** | 0.28901 | < 2e-16 | 0.37742 | < 2e-16 |
| 1.3 | *response ~ trial + EV + **richness*** | 0.50425 | < 2e-16 | 0.87832 | < 2e-16 |
| 2.2 | *response ~ trial + EV + **MB\*5-HT\*richness*** | -0.220810 | 0.108 | -0.36003 | 0.0133 |
| 3.1 | *response ~ trial + EV + **state*** | 3.25338 | <2e-16 | 4.26579 | <2e-16 |
| 3.4 | *state ~ trial + EV + **richness*** | 1.04555 | <2e-16 | 1.21094 | <2e-16 |
| 4.1 | *state ~ trial + EV + **MB\*5-HT*** | -0.25645 | 0.05528 | -0.35446 | 0.00224 |
| 4.2 | *state ~ trial + EV + **MB\*5-HT\*richness*** | -0.99853 | 3.27e-8 | -0.50918 | 0.002849 |
| 4.3 | *pupil size ~ trial + state + **MB\*5-HT*** | -94.95 | 0.10835 | -194.12 | 0.02181 |
| 4.4 | *state increase ~ trial + **pupil\*MB\*5-HT*** | -1.25769 | 0.0258 | -0.64394 | 0.1605 |

## Acknowledgements

The MRS package was developed by Edward J. Auerbach and Małgorzata Marjańska and provided by the University of Minnesota under a C2P agreement.

We thank the Biomedical Services staff at the University of Oxford for expert care of the macaques, and Jenna Ley Greaves, Lucy Underdown, and Sara Rohling for assistance with cannulation procedures.

We are grateful to Luke Priestley for sharing his GLM–HMM analysis scripts and for valuable intellectual input throughout the project.

We thank Tim Viney and Sara Hijazi at the Department of Pharmacology of the University of Oxford, for providing and managing access to the confocal microscope and related software.

We thank Michael Gray and Jake Toth at the Institute of Biomedical Engineering of the University of Oxford for assisting with calibration of the ultrasound transducers.

## Author contributions

- Conceptualization: SW, MFSR, NK
- Methodology: SW, MTN, WR, MH, US, MT, WTC, WWW, DJW, TS, ES, MFSR, NK
- Investigation: SW, MTN, WR, US, NK
- Funding acquisition: MFSR, NK
- Supervision: ES, MFSR, NK
- Writing – original draft: SW, NK
- Writing – review & editing: SW, MTN, WR, MH, US, MT, WTC, WWW, DJW, TS, ES, MFSR, NK

